# Antibiotic hyper-resistance in a class I aminoacyl-tRNA synthetase with altered active site signature motif

**DOI:** 10.1101/2023.01.31.526100

**Authors:** A. Brkic, M. Leibundgut, J. Jablonska, V. Zanki, Z. Car, V. Petrovic-Perokovic, A. Maršavelski, N. Ban, I. Gruic-Sovulj

**Affiliations:** Department of Chemistry, Faculty of Science, University of Zagreb, Horvatovac 102a, 10000 Zagreb, Croatia; Department of Biology, Institute of Molecular Biology and Biophysics, ETH Zürich, 8093 Zürich, Switzerland; Department of Biomolecular Sciences, Weizmann Institute of Science, 7610001 Rehovot, Israel

**Keywords:** isoleucyl-tRNA synthetase, mupirocin, antibiotic resistance, protein synthesis, the HIGH motif

## Abstract

Antibiotics target key biological processes that include protein synthesis. Bacteria respond by developing resistance, which increases rapidly due to antibiotics overuse. Mupirocin, a clinically used natural antibiotic, inhibits isoleucyl-tRNA synthetase (IleRS), an enzyme that links isoleucine to its tRNA^Ile^ for protein synthesis. Two IleRSs, mupirocin-sensitive IleRS1 and resistant IleRS2, coexist in bacteria. The latter may also be found in resistant *Staphylococcus aureus* clinical isolates. Here, we describe the structural basis of mupirocin resistance and unravel a mechanism of hyper-resistance evolved by some IleRS2 proteins. We surprisingly find that an up to 10^3^-fold increase in resistance originates from alteration of the HIGH motif, a signature motif of the class I aminoacyl-tRNA synthetases to which IleRSs belong. The structural analysis demonstrates how an altered HIGH motif could be adopted in IleRS2 but not IleRS1, providing insight into an elegant mechanism for coevolution of the key catalytic motif and associated antibiotic resistance.

## Introduction

Protein synthesis is a central cellular process frequently targeted by natural and man-made antibiotics^1, 2^ that specifically bind to components of the translational machinery. Bacteria, however, frequently develop resistance to antibiotics through mutations or adaptation, making antibiotic resistance an urgent public health problem^3^. Therefore, a better understanding of the resistance mechanisms is of utmost importance.

Aminoacyl-tRNA synthetases (AARSs) play a key role in the fidelity of translation since they catalyze covalent coupling of amino acids to cognate tRNAs^4, 5^. Subsequently, aminoacylated tRNAs (AA-tRNAs) bind to the ribosome in a codon-dependent manner and appropriate amino acids are incorporated into the growing polypeptide chain. AARSs catalyze the formation of AA-tRNA in two steps within the same active site (**Fig. 1A**). The amino acid is first activated by ATP to form an aminoacyl-adenylate (AA-AMP) intermediate, followed by the transfer of the aminoacyl moiety to the tRNA.

**Fig. 1:**
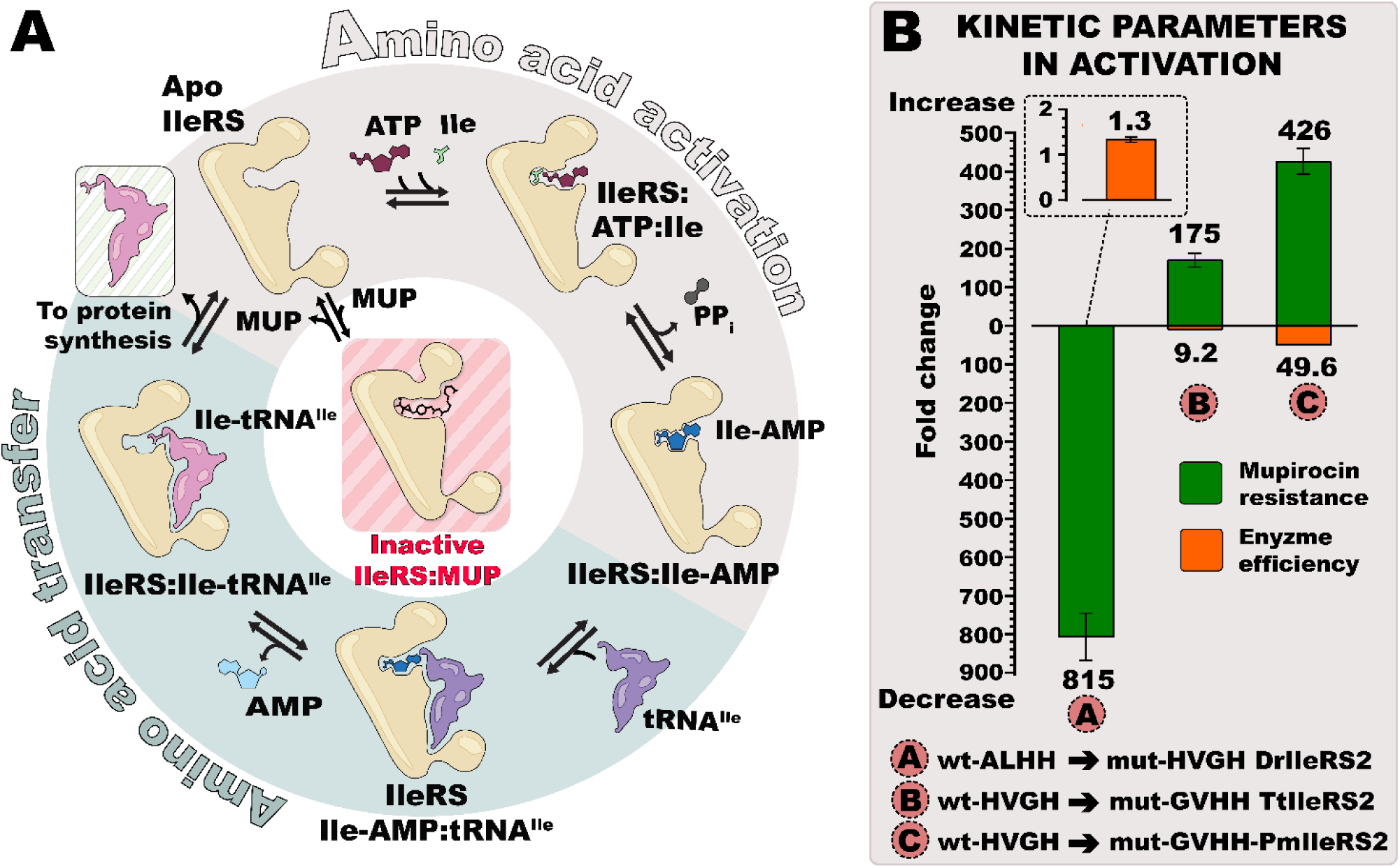
Schematic depiction of the two-step aminoacylation reaction catalyzed by isoleucyl-tRNA synthetase (IleRS). (A) The reaction features formation of a reaction intermediate, isoleucyl-adenylate (Ile- AMP), during the amino acid activation step, after which the isoleucyl moiety is transferred to the 2’-hydroxyl group of the tRNA^Ile^. Bacterial IleRS are susceptible to competitive inhibition by the natural antibiotic mupirocin (MUP)^18^. **(B)** In IleRS2 the non-canonical version (GXHH) of the class I AARS signature motif (HXGH) promotes mupirocin resistance. Exchange of WT non-canonical to canonical (DrIleRS2) or WT canonical to non-canonical (TtIleRS2 and PmIleRS2) signature motifs promotes up to a 10^3^-fold decrease or increase in *K*i for mupirocin, respectively, measured at the activation step (green columns). At the same time, the catalytic efficiency (*k*cat/*K*M) of the mutants in the activation step was only slightly compromised (orange columns). “X” in HXGH and G/AXHH indicates a variable position and stands for Ile/Val/Leu/Met/Tyr.

AARSs are divided into two Classes, I and II, each characterized by class-dependent catalytic folds^6^ and sequence motifs^7^. Class I AARSs share the nucleotide-binding fold with Rossmann-like topology that belongs to the larger HUP superfamily^8^ and two, so-called, signature motifs, the HIGH and KMSKS motifs. The HIGH motif is located at the tip of helix α1 of the conserved catalytic core comprising segments β1-α1-β2-α2-β3 and α3-β4-α4-β5 that are separated by peptide insertions CP 1 and 2 (**Supplementary Fig.1)**. The KMSKS sequence is located on the flexible loop that follows the β5 strand. Both motifs, highly conserved in Class I AARSs^9^, are an integral part of the active site and are essential for ATP binding and stabilization of the transition state for amino acid activation^10–12^.

Isoleucyl-tRNA synthetase (IleRS) is a Class I AARS inhibited by the clinically used antibiotic mupirocin (commercial name Bactroban^®^) that is naturally produced by *Pseudomonas fluorescences* ^13^, which competes with isoleucine and ATP for binding at the active site^14, 15^. However, two types of IleRSs exist in bacteria that, apart from displaying distinct sequences for the C-terminal tRNA anticodon binding domains, also feature different susceptibility of the active site to mupirocin^16, 17^. Specifically, type 1 proteins (IleRS1) are strongly inhibited by mupirocin (*K*i in low nanomolar range^18, 19^), while IleRS2, which are homologous to IleRSs from the eukaryote cytosol^16, 17^, exhibit resistance to mupirocin concentrations that are about three orders of magnitude higher^20^. In some cases, they even reach millimolar levels^17^, which we term here “hyper-resistance”. IleRS1 and IleRS2 generally occur individually in bacteria, but there is a small group of *Bacillaceae* that carry both genes in the genome^21^. IleRS2 plays an important role in providing mupirocin resistance to bacteria in hospital environments, since mupirocin-resistant *Staphylococcus aureus* isolates acquired the *ileS2* gene on a plasmid^22^. Comparison of the crystal structures of *S. aureus* IleRS1 bound to mupirocin and tRNA^14^ and *Thermus thermophilus* IleRS2 bound to mupirocin only^15^, provides some indications of why in type 2 IleRS the affinity for mupirocin is reduced. However, the comparison is difficult since the structures include different ligands.

Using a combination of phylogenetic, biochemical and X-ray structural analysis we investigated the basis for mupirocin resistance and the origin of hyper-resistance in IleRS2. We found that some IleRS2 harbor an altered Class I HXGH signature motif (with X representing a hydrophobic residue) such that the 1^st^ and the 3^rd^ amino acids are swapped. This GXHH altered signature motif conveys IleRS2 with hyper-resistance to mupirocin, while catalytic activity is only mildly affected. We determined structures from *Priestia (Bacillus) megaterium*^23^ wild-type IleRS1 and IleRS2, both carrying the canonical signature motif, as well as mutants with a swapped GXHH motif, complexed to an aminoacyl-adenylate analog or mupirocin. These findings revealed why the altered HXGH motif could not be introduced in IleRS1 without abolishing catalysis. Our results provide new insights into the mechanism of antibiotic hyper-resistance and evolution of tRNA synthetases under selective pressure.

## Results

### Mupirocin hyper-resistance in type 2 IleRS is related to the altered class I signature motif

To investigate whether sequence analysis may shed light on the origin of hyper-resistance occurring in some type 2 IleRSs, the sequences of 379 IleRSs were retrieved from representative prokaryotic proteomes^24^, aligned, and the phylogenetic tree was inferred. We found that IleRS2 grouped in two distinct clades (**Fig. 2A, Supplementary Data 1)**, each, surprisingly, containing a number of IleRS2 lacking the canonical signature HIGH motif (we refer to it as HXGH, as various hydrophobic amino acids are found at the 2^nd^ position, **Fig. 2B**, **Supplementary Data 2)**. Instead, their motifs (dubbed non-canonical) have His at the 3^rd^ position, while the 1^st^ position is occupied by Gly or rarely Ala. Therefore, the newly found non-canonical motif (G/AXHH), in essence, has the 1^st^ and the 3^rd^ position swapped. The exchange is not trivial; the 1^st^ His stabilizes the transition state of the amino acid activation step, and its mutation in several class I AARSs decreases the corresponding rate by a 10^3^- fold^10, 12^, while Gly is strongly conserved to allow accommodation of the adenine base. We realized that IleRS2 from *Streptomyces griseus* (SgIleRS2), which exhibits a *K*i for mupirocin in mM range^17^, harbors the same non-canonical motif. Thus, we wondered whether this highly unexpected change in the class I signature motif could be related to mupirocin hyper- resistance.

**Fig. 2:**
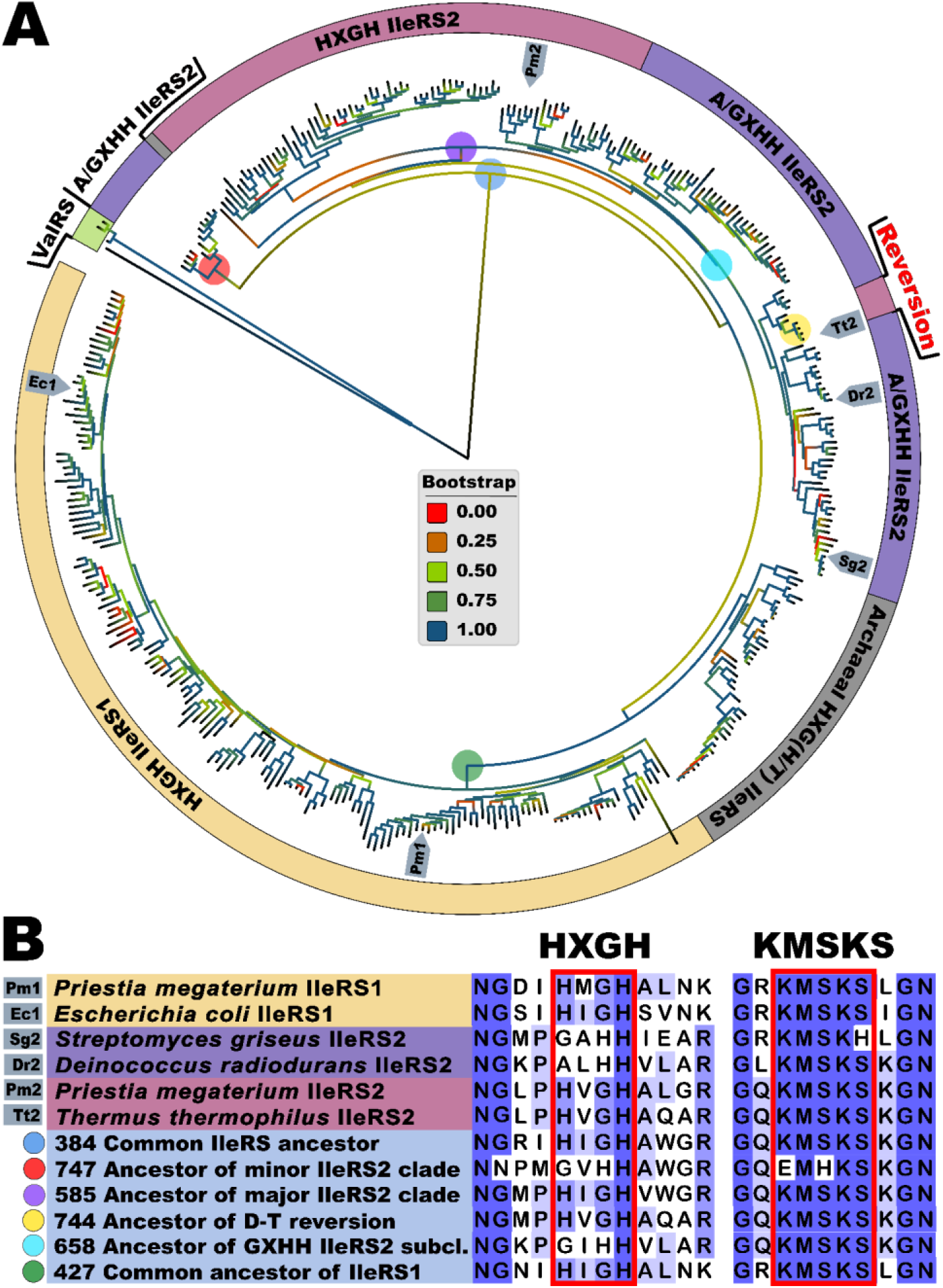
Phylogenetic analysis of prokaryotic IleRSs. (A) The tree was constructed from a multiple sequence alignment of the enzyme’s PFAM domains using 379 IleRS sequences (158 HXGH IleRS1, 41 archeal IleRS, 77 HXGH IleRS2, and 86 G/AXHH IleRS2). The tree and the alignment are given in **Supplementary Data 1** and **2**. The HXGH motif is present in all IleRS1. IleRS2 can accommodate either HXGH or non-canonical G/AXHH. Naturally occurring reversion of the non-canonical to the canonical motif occurred in the *Deinococcus-Thermus* clade. ValRS sequences were used as an outgroup. A low bootstrap of the IleRS2 early branch is likely a consequence of a small number of available sequences from *Plantomycetota* and *Chlorflexi* phyla. The IleRS enzmyes used in this study are marked on the tree. **(B)** Sequence alignment of the HXGH and KMSKS signature motifs for selected IleRS1 and IleRS2 enzymes and the key ancestral nodes in IleRS evolution. The numbers describe the position of the node in the ancestral tree presented at **Supplementary** Fig. 4 **and Supplementary Data 3**. The sequences of the nodes are given in **Supplementary Data 4**.

To explore the mechanism of (hyper-) resistance in greater depth, we tested the mupirocin susceptibility of *Deinococcus radiodurans* IleRS2 (DrIleRS2), which also carries a non-canonical motif naturally, in this case ALHH. A classical competitive inhibition with respect to both Ile and ATP was observed in the amino acid activation step (**Table 1, Supplementary Table 1, Supplementary Fig. 2)**. The measured *K*i was 6.6 mM, which is more than a 10^3^-fold higher than the *K*i of the IleRS2 enzymes carrying the canonical HXGH motif (**Table 1**) and similar to the SgIleRS2 with the non-canonical motif^17^. Next, we exchanged the natural non-canonical motif of DrIleRS2 with the canonical (ALHH with HVGH). The mutant, mut-HVGH-DrIleRS2, exhibited an 815-fold drop in *K*i, corresponding to a drastic loss of resistance, thus linking the non-canonical motif directly with hyper-resistance (**Fig. 1B**, **Table 1**). We questioned whether the motif exchange (i.e., the swap of the 1^st^ and the 3^rd^ motif position) will increase the resistance of the canonical IleRS2. To address this, we chose two IleRSs from species where the canonical HXGH is present, *T. thermophilus* IleRS2 (TtIleRS2) and *P. megaterium* IleRS2 (PmIleRS2), and exchanged their canonical HVGH motif with GVHH. Both mutants (mut-GVHH-TtIleRS2 and mut-GVHH- PmIleRS2) indeed experienced a 200-fold increase in *K*i (**Fig. 1B**, **Table 1**). Altogether, our data provide compelling evidence that the non-canonical form (G/AXHH) of the Class I AARS signature motif is responsible for up to a 10^3^-fold increase in mupirocin resistance in IleRS2.

**Table 1.** Steady-state amino acid activation parameters and mupirocin inhibition constants^a^

| | Signature motif | substrate | $k_{\text{cat}} / \text{s}^{-1}$ | $K_{\text{M}} / \mu\text{M}$ | $K_{\text{i}} (\text{MUP})_{\text{Ile}} / \mu\text{M}$ |
| --- | --- | --- | --- | --- | --- |
| wt-ALHH<br>DrIleRS2 | Natural<br>non-canonical | Ile | $45.1 \pm 0.3$ | $44 \pm 1$ | $6600 \pm 400$ |
| | | ATP | | $580 \pm 20$ | |
| mut-HVGH<br>DrIleRS2 | Mutated to<br>canonical | Ile | $24.3 \pm 0.3$ | $17.8 \pm 0.5$ | $8.1 \pm 0.5$ |
| | | ATP | | $390 \pm 2$ | |
| wt-HVGH<br>TtIleRS2 | Natural<br>canonical | Ile | $27.9 \pm 0.3$ | $18.7 \pm 0.6$ | $0.20 \pm 0.01$ |
| | | ATP | | $2200 \pm 100$ | |
| mut-GVHH<br>TtIleRS2 | Mutated to<br>non-canonical | Ile | $6.8 \pm 0.2$ | $42 \pm 2$ | $34 \pm 2$ |
| | | ATP | | $3300 \pm 100$ | |
| wt-HVGH<br>PmIleRS2<sup>#</sup> | Natural<br>canonical | Ile | $66 \pm 2$ | $49 \pm 2$ | $1.08 \pm 0.06$ |
| | | ATP | | $1600 \pm 100$ | |
| mut-GVHH<br>PmIleRS2 | Mutated to<br>non-canonical | Ile | $6.4 \pm 0.1$ | $237 \pm 5$ | $460 \pm 10$ |
| | | ATP | | $3400 \pm 100$ | |
| wt-HMGH<br>PmIleRS1<sup>#,b</sup> | Natural<br>canonical | Ile | $30 \pm 2$ | $2.1 \pm 0.3$ | $0.00029 \pm 0.00002$ |
| | | ATP | | $395 \pm 32$ | |

<sup>a</sup> Measured using ATP/PP<sub>i</sub> exchange assay, source data are given in Supplementary Data 5<sup>b</sup> Slow tight binding inhibitor<sup>#</sup> Values from<sup>21</sup>The values represent the average value ± SEM of at least three independent experiments

### The non-canonical class I signature motif cannot be functionally accommodated in IleRS1

Alteration of the HXGH motif in some IleRS2 was highly surprising considering the key catalytic role of this motif in the amino acid activation. Hence, we explored how this is accomplished, and whether it comes at a trade-off with IleRS activity, using kinetics and structural approaches.

As shown in Table 1, DrIleRS2, which naturally carries the non-canonical motif, shares catalytic efficiency in the activation step (*k*cat/*K*M) with the canonical PmIleRS2 and TtIleRS2, supporting a lack of catalytic trade-offs related to hyper-resistance. Introducing the non-canonical GVHH motif into PmIleRS2 and TtIleRS2 also did not strongly affect the enzymes, yet hyper-resistance, in this case, came at the expense of increased *K*M and decreased *k*cat values of up to 10-fold (**Table 1**, **Fig. 1B**). Finding that the non-canonical motif is well tolerated in IleRS2 is consistent with its broad distribution among IleRS2s (**Fig. 2**).

In sharp contrast, the phylogenetic analysis did not identify a single IleRS1 with the non-canonical motif (**Fig. 2**), strongly suggesting that HXGH motif variations are not tolerated among IleRS1. To test this, we exchanged the natural HXGH motif in IleRS1 from *P. megaterium* and *Escherichia coli* with GXHH. GXHH-mutants showed a lack of product formation during prolonged reaction times even at 15 µM enzyme (**Supplementary Fig. 3**), confirming that the active site of IleRS1 cannot productively accommodate the non-canonical motif.

To reconstruct the evolutionary origin of the non-canonical motif, we inferred the IleRS ancestral states from our phylogenetic analysis (**Fig. 2B, Supplementary Fig. 4, Supplementary Data 3 and 4**). The HXGH motif is found in most of the earliest ancestors, i.e., the common ancestor of all IleRSs (node 384), the IleRS1 ancestor (node 427) and the ancestor of the major IleRS2 clade (node 585). The exception is the ancestor of the minor IleRS2 clade (node 747), which harbors the non-canonical GXHH motif. Among the inner nodes, GXHH can be found at node 658, which represents the GXHH subclade in the major IleRS2 clade. The presence of GXHH in two distant IleRS2 nodes indicates that the non-canonical motif was acquired at least twice during evolution of IleRS2. That the motif exchange is not detrimental to IleRS2 is also supported by a natural reversion of the non-canonical to the canonical motif in the *Deinococcus-Thermus* clade (node 744, ancestor has the HXGH motif). That said, laboratory exchange of HVGH back to GVHH in TtIleRS came at a minor expense of its catalytic efficiency (Table 1). The observation that all early ancestors (except node 747) carry the class I AARS-dominating HXGH motif may indicate that both IleRS2 and IleRS1 emerged with the HXGH motif. Mutation of the IleRS2 signature motif appears later in the evolution, presumably under selective pressure to withstand higher antibiotic concentration in the environment.

### Structural basis of the non-canonical motif accommodation solely in IleRS2

To understand how IleRS2, but not IleRS1, accommodates the non-canonical motif, we used X-ray crystallography to determine the structures of a pair of IleRS enzymes from the same organism. Apart from wild-type PmIleRS1 and PmIleRS2, both of which harbor the canonical HXGH motif, we also structurally characterized their corresponding GXHH mutants, all in complex with a non-hydrolyzable analogue of the isoleucyl-adenylate reaction intermediate, Ile-AMS (**Supplementary Table 2, Supplementary Fig. 5**).

The overall fold of the full-length wild-type PmIleRS1 and PmIleRS2 (**Figs 3A and 3B**) and the topology and architecture of their active sites (**Supplementary Fig. 1**) correspond well to the previously determined structures of IleRS1 from *S. aureus*^14^ and IleRS2 from *T. thermophilus*, *Candida albicans and Saccharomyces cerevisiae*^15, 25, 26^. Additionally, for the first time, the structure of the C-terminal tRNA anticodon binding domain, which differs among the two IleRS types^16^, is resolved for a type 2 protein (**Fig. 3A and Supplementary Fig. 6**). The structures reveal that the reaction intermediate analogue Ile- AMS in both wild-type enzymes binds to the active site in a canonical manner^15^ (**Supplementary Fig. 7**). However, superimposing the HUP catalytic cores of IleRS1 (residues 50-174 and 523-635) and IleRS2 (residues 40-168 and 513-632) unraveled conformational rearrangements within the signature motifs in the active sites (**Fig. 3**). Relative to IleRS1, the KMSKS loop assumed a more closed conformation in IleRS2, as indicated by a 5.0 Å displacement of the 1^st^ Lys backbone towards helix α1 (**Fig. 3C, left**). While the KMSKS loop is known to be flexible and to change its conformation during the reaction^27^, we also found that the HXGH motif in IleRS2 is displaced towards the KMSKS loop (**Fig. 3C**). The latter affects repositioning of the 1^st^ and the 4^th^ histidine of the HXGH motif by 2.3 Å and 2.4 Å, respectively, and the Gly α-carbon by 1.7 Å relative to the analogous positions in IleRS1 (**Fig. 3C, right**). The displacement (shift) of the HIGH motif in IleRS2 is of utmost importance for the accommodation of the non-canonical A/GXHH motif and emergence of hyper-resistance, as discussed below. Hence, to explore it further, we extensively analyzed the residues in the immediate vicinity of the HXGH motif using mutagenesis combined with biochemical and structural characterization (exemplified for the W130Q PmIleRS2 mutant in **Supplementary Fig. 8 and Supplementary Table 2**). However, no simple mechanism responsible for the observed conformational change that remodels the IleRS2 active site emerged, suggesting that the shift of the HXGH motif is more complex and likely deeply rooted in the architecture of the IleRS2 active site.

**Fig. 3:**
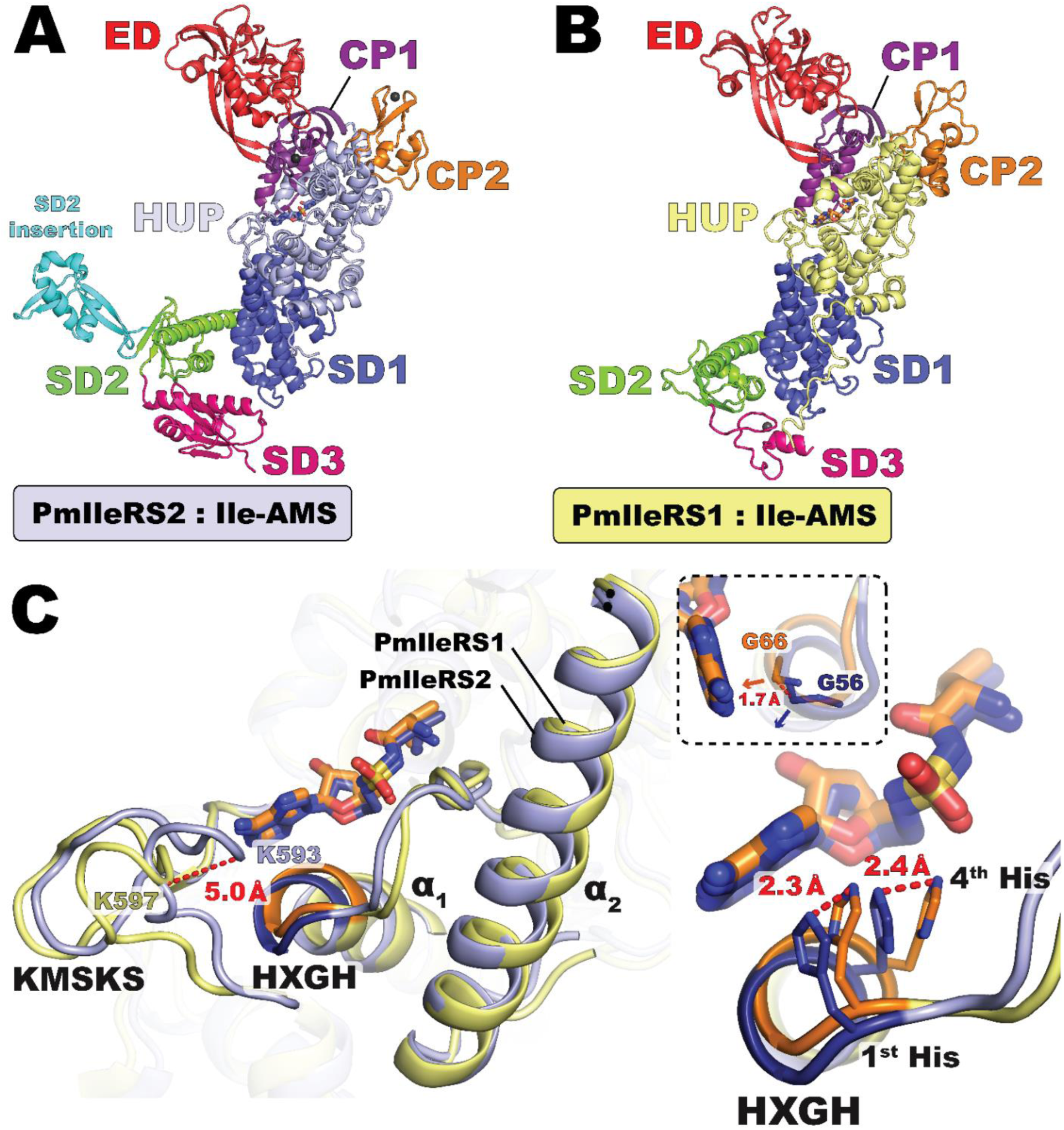
**Comparison of crystal structures of PmIleRS2 and PmIleRS1 in complex with the non-hydrolysable analogue of the reaction intermediate Ile-AMS**. **(A, B)** The canonical structures with the HUP catalytic domain, the CP1 and CP2 domains (CP refers to connective peptide) and the editing domains (ED) inserted into CP1 are visualized. For the first time, the full-length C-terminal domain of type 2 IleRS is resolved, revealing three subdomains (SD, **Supplementary** Fig. 6), among which SD3 differs in size, fold and lack of the zinc-binding motif relative to SD3 in IleRS1. A novel insertion into SD2 is observed in IleRS2. **(C)** Structural overlay of the IleRS1 and IleRS2 HUP catalytic cores (residues 50-174 and 523-635 in IleRS1 and 40-168 and 513-632 in IleRS2) bound to Ile-AMS revealed overlapping positions of Ile-AMS and a conformational rearrangement of the active site in IleRS2 relative to IleRS1. The tip of helix α1 comprising the HXGH motif (blue in IleRS2 and orange in IleRS1) and the KMSKS loop move towards each other in IleRS2. This repositions both the 1^st^ and 4^th^ histidine residues (right panel) as well as the glycine α-carbon (inset).

How does alteration of the canonical motif (i.e., exchange of the 1^st^ and the 3^rd^ position in HXGH) influence the PmIleRS structures and, hence, their function? At the level of the overall fold, the WT enzymes and corresponding GXHH mutants are highly superimposable (**Supplementary Figs 9A and 9D**). However, while the binding of Ile-AMS to the active sites of both GXHH mutants parallels its binding to the corresponding wild-type enzymes (**Fig. 4**, **Supplementary Fig. 7)**, some distinctions that are mainly related to the precise accommodation of the adenine base emerged. In PmIleRS1, introducing the GMHH mutation leads to a 1.5 Å shift of the adenine base and a 2.1 Å move of Phe586, which loses its stacking with the adenine moiety (**Fig. 4A**). Such a distorted geometry of the active site likely results in a non-productive binding of the ATP substrate and concomitant loss of mut- GMHH-IleRS1 activity (**Supplementary Fig. 3**). Such a scenario does not occur in PmIleRS2, where the tip of the α1 helix in both the mut-GVHH-PmIleRS2 and corresponding wild-type enzyme are similarly rearranged (shifted) compared to IleRS1, resulting in a 1.7 Å displacement of the backbone in the third position of the motif (**Fig. 3C, right**). This allows accommodation of the 3^rd^ His in mut-GVHH-PmIleRS2 without displacement of the adenine moiety. This way, the 3^rd^ His can take over the role of the 1^st^ His in the canonical reaction mechanism (**Fig. 4B**).

**Fig. 4:**
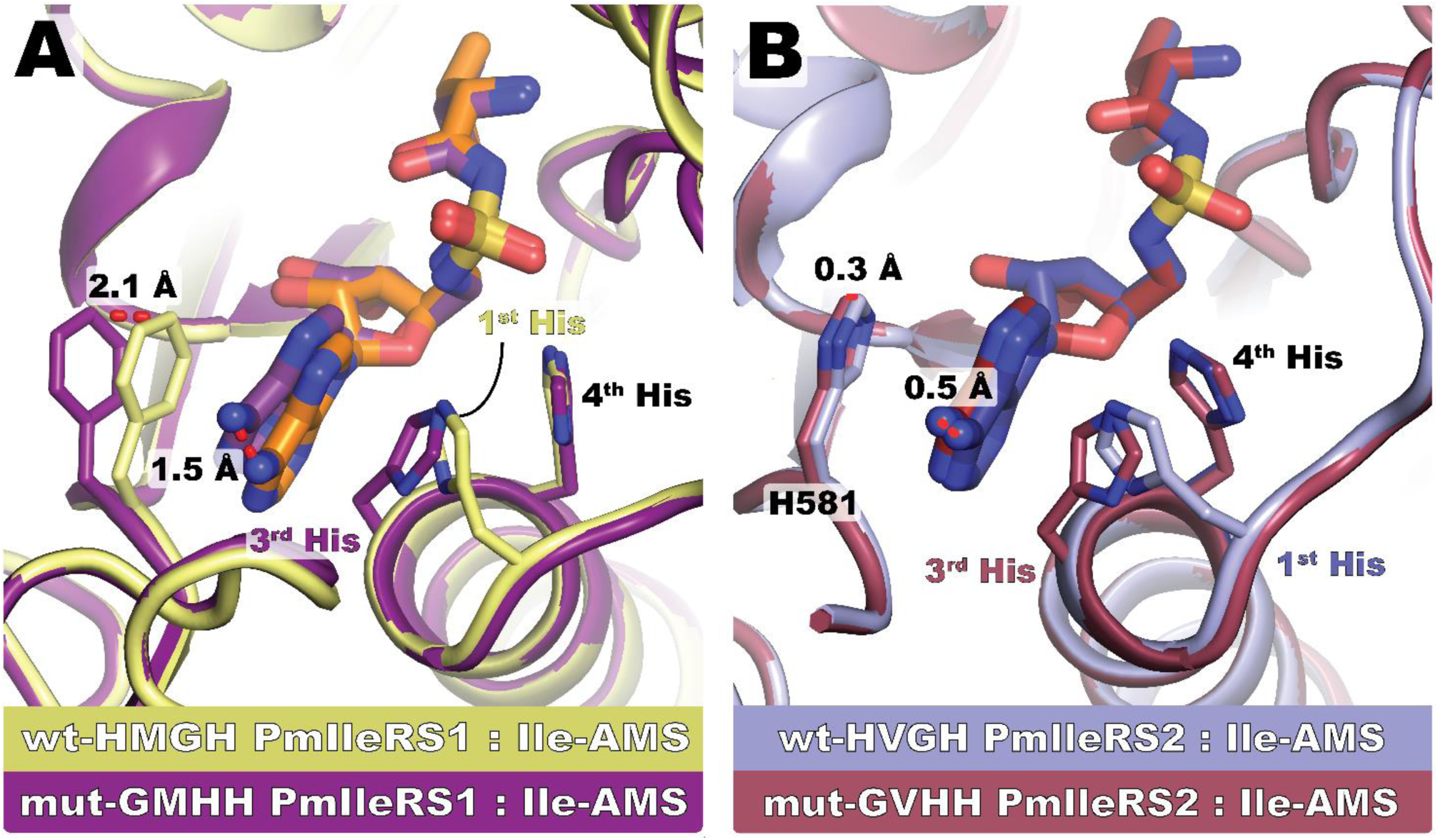
Structural overlay of the active sites of WT PmIleRS1 and PmIleRS2 with mutants where histidines in the signature motif are exchanged. (A) In IleRS1, the 3^rd^ His from GV<u>H</u>H promotes mispositioning of the adenine base that contributes to the abolished catalysis. **(B)** In IleRS2, the general shift of the GMHH motif relative to IleRS1 allows accommodation of a His in the 3^rd^ position without mispositioning of the adenine moiety. In both the mut and wt-structures, the 3^rd^ His HNɛ2 from GXHH adopts an equivalent position as the catalytically relevant HNɛ2 of the 1^st^ His from HXGH.

### Structural basis of mupirocin resistance

The non-canonical motif endows IleRS2 with hyper-resistance, the structural basis of which cannot be assessed by crystallography due to a *K*i in the mM range. Therefore, to deepen our understanding of mupirocin resistance in general and to infer the mechanism of hyper-resistance, we determined the crystal structures of wild-type PmIleRS1 (sensitive) and PmIleRS2 (resistant), both of which carry the canonical HXGH motif and are amenable in mupirocin-bound form when the inhibitor is supplied at sufficiently high concentrations (**Fig. 5, Supplementary Table 2, Supplementary Fig. 10**). The overall structures of mupirocin- bound PmIleRS1 and PmIleRS2 overlap well with those determined in complex with Ile-AMS (**Supplementary Figs 9C and 9E**). In both enzymes, mupirocin binds to the active site mimicking the interaction intermediate Ile-AMP (**Supplementary Figs 9C and 9E**), as expected based on the previous structural data^14, 15^ and the competitive mode of inhibition (**Table 1**). The binding of mupirocin to both (sensitive and resistant) enzymes (**Supplementary Fig. 11**) is similar as described^15^, however we are able to reveal novel interactions. We observe the backbone of Pro56 and the side chain of Asn70 forming interactions with O13 and O7 atoms of mupirocin, respectively, in PmIleRS1, which are absent in PmIleRS2 (**Figs 5B and 5C, middle**). Furthermore, stacking interactions of Phe586 and the unsaturated C2-C3 bond of the monic acid A part (**Fig. 5A**) contributes to the affinity in IleRS1, while the analogous His582 is reoriented in PmIleRS2 (**Figs 5B and 5C, middle**). Finally, we found that the carboxylate oxygen of the nonanoic acid part of mupirocin (**Fig. 5A**) establishes an H-bond with the backbone NH of K597 from the KMS<u>K</u>S loop only in PmIleRS1 (**Figs 5B and 5C, middle**). It was demonstrated^28, 29^ that H-bonding to a charged group may contribute 3-4 kcal/mol to binding, which translates into up to a 10^3^ contribution to a binding constant. Indeed, H-bonding of the tyrosyl-tRNA synthetase active site aspartate and the hydroxyl group of the tyrosine substrate is estimated to contribute a 10^3^-fold to the selectivity of the enzyme against phenylalanine^28^. In PmIleRS2, the closed conformation of the KMSKS loop (**Fig. 5C, middle**) forces binding of the nonanoic carboxylate such that it is positioned at 4.1 Å distance to the NH of Lys593 (KMS<u>K</u>S). It is this lack of carboxylate stabilization by H-bonding that may strongly affect the IleRS2 affinity towards mupirocin and provide resistance.

**Fig. 5:**
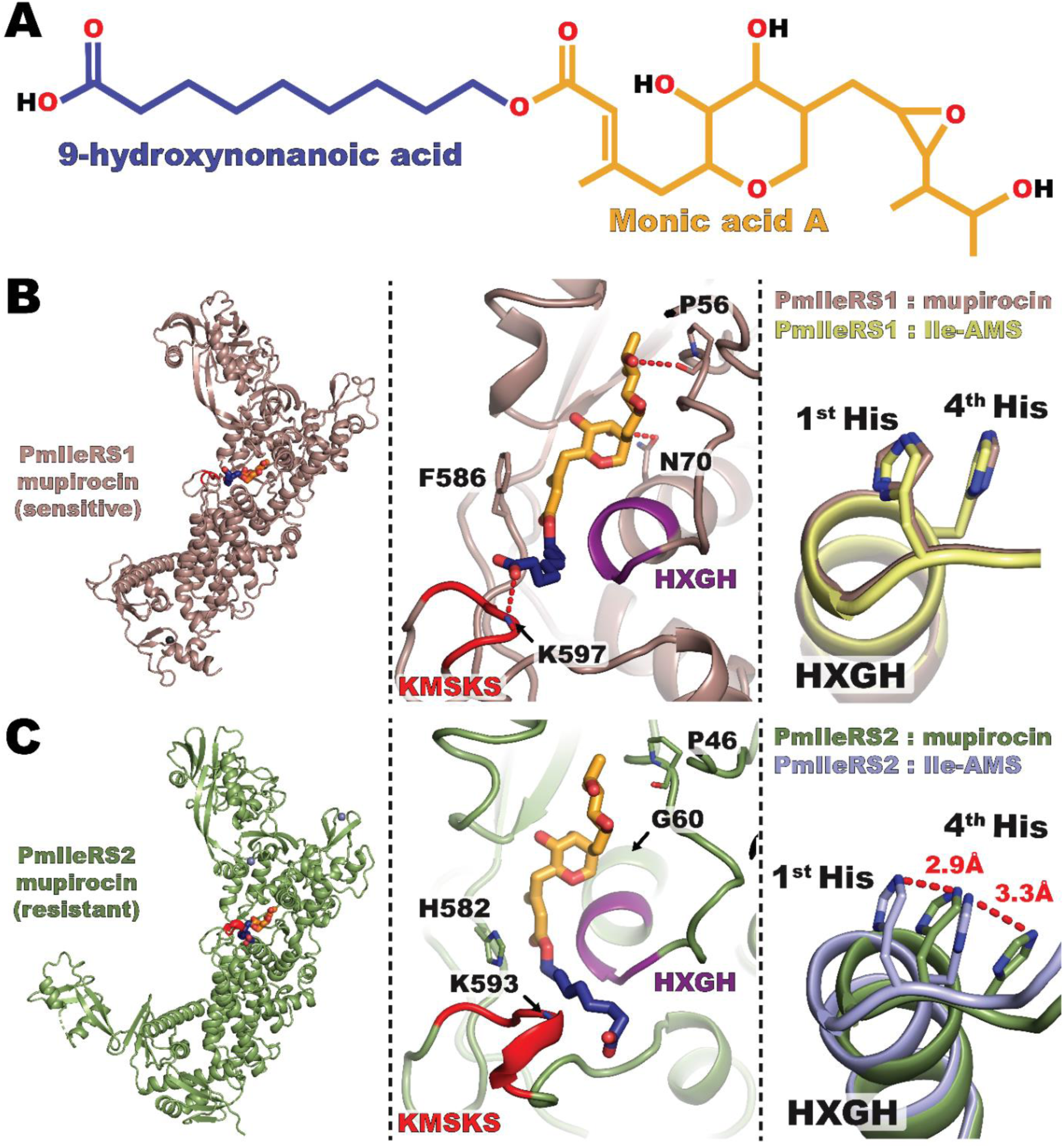
Structures of PmIleRS1 and PmIleRS2 bound to mupirocin. (A) Mupirocin is an ester of monic acid A and 9-hydroxynonanoic acid. Within the active sites of IleRS1 and IleRS2, the monic acid part mimics the Ile-AMP interactions, whilst the nonanoic acid part is oriented towards the KMSKS loop. **(B)** In IleRS1, unique interactions with mupirocin include the carboxyl group of the nonanoic acid moiety, which establishes an H-bond with the backbone NH of K597 in the KMSKS loop, the hydrogen bonds between the monic acid part and a proline and an asparagine and stacking interactions of phenylalanine. The position of the HXGH motif remains unaltered upon mupirocin binding. All interactions are depicted in **Supplementary** Fig. 11. **(C)** In the IleRS2 active site, mupirocin binds via a lower number of interactions, providing the basis for resistance. Stabilization of the nonanoic carboxylate by H-bonding is precluded by a closed conformation of the KMSKS loop, thereby forcing the nonanoic acid moiety into the cleft between the HXGH motif and the KMSKS loop. As a result, the HXGH motif is pushed away from the KMSKS loop.

Interestingly, we observed a kink of helix α2 (from residues 123–126) solely in the crystal structure of mupirocin-bound PmIleRS2 (**Supplementary Fig. 12A**). To explore whether these structural changes in PmIleRS2 are due to different conformational readjustments of the active site upon ligand binding (i.e., induced fit)^30^ or sampling of different enzyme’s conformations from solution (i.e., conformational selection)^31^, we aimed to crystallize IleRS2 in the apo form. Because this approach remained unsuccessful, we used molecular dynamics (MD) to address the conformation of the helix α2 in the absence of ligands. When we removed the ligands from the PmIleRS2 structures, we found that independent of whether we started from the apo structure with the regular helix α2 (obtained by removing Ile-AMS) or the distorted one (obtained by removing mupirocin), helix α2 predominately (>99%) went into the distorted conformation during 360 ns simulations (**Supplementary Fig. 12B)**. Noteworthy, an irregular helix α2 structure was also observed in MD simulations of IleRS2 bound to Ile-AMS, yet here distortion was observed after around 180 ns (**Supplementary Fig. 13**). Thus, the MD data suggest that the kink in helix α2 is part of the conformational flexibility of IleRS2 and is neither induced by mupirocin binding nor by different packing environments in the different space groups that PmIleRS2:mupirocin and PmIleRS2:Ile-AMS crystallized in (**Supplementary Table 2**). Instead, it appears that Ile- AMS and mupirocin sample different IleRS2 conformations that exist in solutions.

### Modelling of hyper-resistance in PmIleRS2

Structural data unraveled that in the PmIleRS2:mupirocin complex, the tip of the helix α1 carrying the HXGH motif is shifted back towards helix α2, resulting in 2.9 Å or 3.3 Å displacements of the 1^st^ or 4^th^ His, respectively, relative to PmIleRS2:Ile-AMS (**Fig. 5C, right**). No such ligand-dependent rearrangements in the HXGH motif are observed in PmIleRS1 (**Fig. 5B, right**). Thus, in the PmIleRS2:mupirocin complex the HXGH motif adopts the same position as in the IleRS1 structures (**Supplementary Fig. 14A)**. The back- shift of the HIGH motif is likely promoted by binding of the nonanoic part of mupirocin to the channel between the HIGH motif and the KMSKS loop forced by the closed conformation of the KMSKS loop in PmIleRS2. (**Supplementary Fig. 14B**).

We hypothesize that this back-shift, which places the 3^rd^ amino acid of the HXGH motif into close proximity to the adenine base, provides the basis of A/GXHH motif-promoted hyper-resistance. To address this hypothesis, we *in silico* exchanged the canonical motif in the structure of wt-HVGH-PmIleRS2 bound to mupirocin with the GVHH sequence. The coordinates for the latter were taken from the mut-GVHH-PmIleRS2:Ile-AMS structure. The resulting clash due to a His in the 3^rd^ position with both, the nonanoic part and pyrrole ring of mupirocin **(Fig. 6)**, likely explains how the presence of the non-canonical motif prevents mupirocin from binding and leads to the further substantial increase in *K*i.

**Fig. 6:**
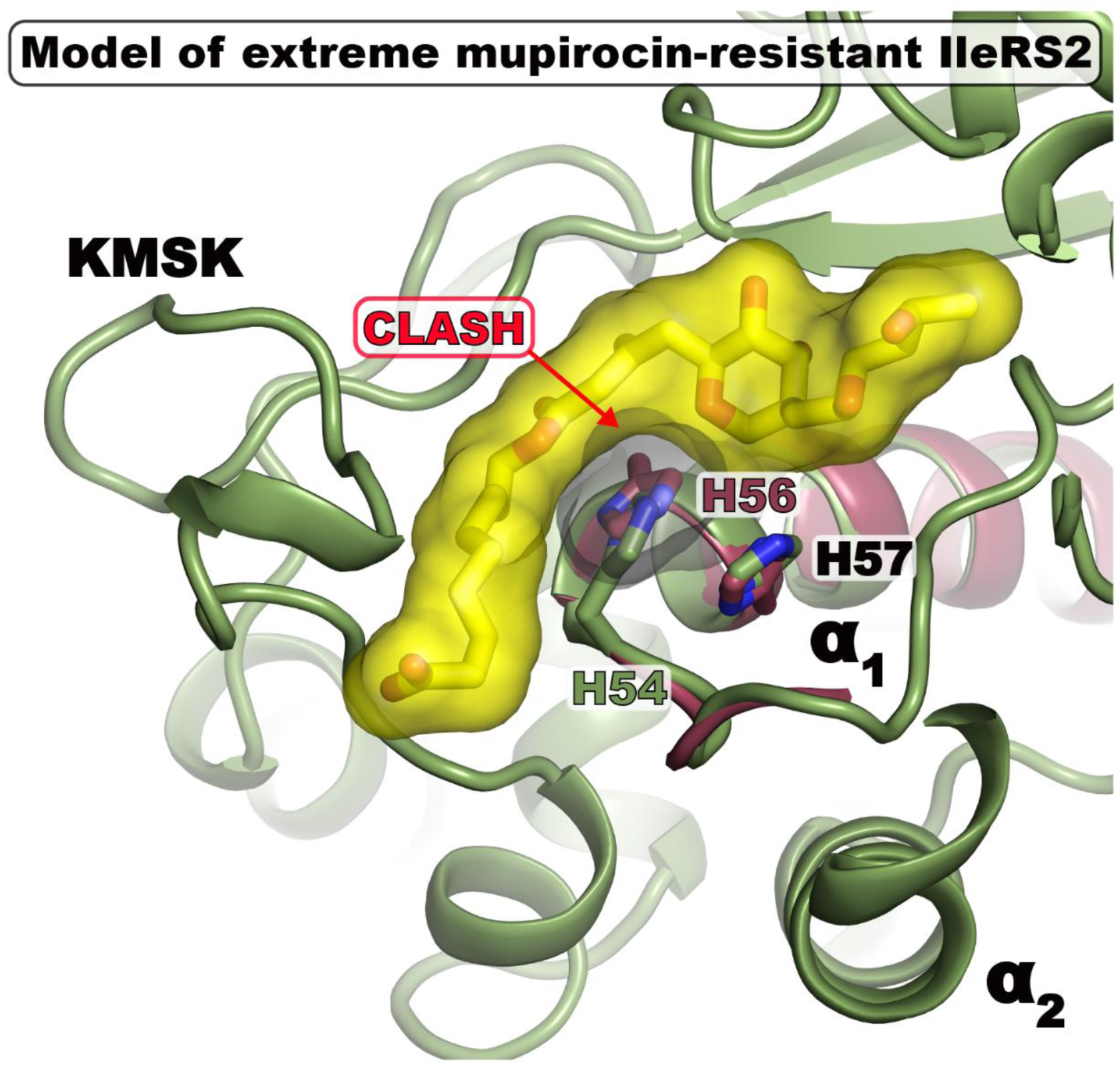
Mupirocin modelled into the active site of mut-GVHH-IleRS2 displaying a hyper-resistance phenotype. The coordinates of the GVHH motif were taken from the mut-GVHH-PmIleRS2:Ile-AMS structure and were superimposed onto wt-HVGH-PmIleRS2:mupirocin. A G56H exchange results in clashes with distances below 2.5 Å excluding hydrogens, consistent with the hyper-resistance observed in IleRS2 variants harboring the non-canonical motif (**Table 1**).

## Discussion

Translation is a central cellular process and a frequent target for antibiotic action. Most of the known translation-related antibiotics act on ribosomes^32^. A smaller number target AARSs, the essential enzymes that secure correctly aminoacylated tRNAs.^2^ Among them, the best-known is mupirocin, which blocks IleRS by binding to both the isoleucine and ATP binding pockets^15^ and is primarily active against gram-positive pathogens, including methicillin-resistant *S. aureus* (MRSA).^33, 34^

In bacteria, two IleRS types, IleRS1 and IleRS2, may perform the housekeeping function. Mupirocin strongly inhibits IleRS1 and modestly IleRS2, with the difference in *K*i ranging from 10^3^-10^5^-fold. Accordingly, the presence of IleRS2 as a sole^17^ or a second^21^ gene in the genome or on a plasmid^22^ confers the resistance. Antibiotic resistance is a major public health problem worldwide^3^ and resolving the underlying mechanisms by which it evolves and operates broadens our capacity to tackle it.

The mechanism of mupirocin resistance has not yet been fully understood. The structural basis was indecisive because comparison of the mupirocin-bound structures of *T. thermophilus* IleRS2^15^ and *S. aureus* IleRS1^14^ is not straightforward considering that the IleRS1 structure besides mupirocin also contains the tRNA, which may promote conformational changes of the active site loops. To alleviate this problem, we solved the structures of IleRS1 (sensitive) and IleRS2 (resistant) from *P. megaterium* bound to mupirocin only (**Fig. 5**). The structures provide an important progress in understanding the resistance mechanism by identifying that the closed conformation of the active site KMSKS loop in IleRS2 precludes H-bonding stabilization of the mupirocin’s carboxylate (**Fig. 5B**, **Supplementary Fig. 14B**). Because H-bonding to a charged group has been shown as a powerful mechanism to ensure a high level of specificity in molecular recognition^29^ and may strongly contribute to binding of mupirocin to IleRS1, loss of this interaction in IleRS2 may have been adopted as an efficient resistance mechanism during evolution. In accordance, no H-bonding to the mupirocin’s carboxylate was observed in the TtIleRS2:mupirocin structure^15^ or in a model of mupirocin bound to eukaryotic IleRS2^25^. The structural data are also consistent with the finding that monic acid A alone (**Fig. 5A**) cannot act as an inhibitor^35^. A possible role of tRNA in modulating the binding of mupirocin seems unlikely, as kinetic data showed that the *K*i for mupirocin in the amino acid activation and the two-step tRNA aminoacylation reactions are similar^17, 18, 36^. Consistent with these findings, no direct interaction between mupirocin and the tRNA was observed in the structure of SaIleRS1 complexed to tRNA and mupirocin (PDB 1QU2)^14^. Further, the active site residues contacting mupirocin in our PmIleRS1:mupirocin structure (**Fig. 5B**) and the SaIleRS1:tRNA:mupirocin complex, including the KMSKS loop and the tip of the α1-helix harbouring the HIGH motif, adopt a highly similar arrangement, indicating that their conformation is independent of the presence of tRNA. Nevertheless, based on our structural findings we cannot exclude that upon the binding of the 3’-end of the tRNA to the catalytic site, the KMSKS loop, which displays increased local temperature factors, may become directly or indirectly stabilized or change its conformation.^37^

A major surprise, however, came from the finding that hyper-resistance (*K*i in mM range) found in some IleRS2 (Table 1 and ^17^) originate from their naturally altered key catalytic motif, the HXGH motif. We mimic the motif alteration by the exchange of the 1^st^ and 3^rd^ motif positions in PmIleRS2 that naturally carries the canonical HVGH motif. This GVHH- PmIleRS2 mutant with a bulky histidine, instead of glycine, at the 3^rd^ position displayed a 460-fold increase in *K*i for mupirocin (**Table 1**) due to a severe steric clash of the 3^rd^ histidine with mupirocin (**Fig. 6**). Motif exchanges in both directions (canonical to non-canonical and *vice versa*) that we carried out in several IleRS2 enzymes unambiguously link the non-canonical motif with hyper-resistance.

Mutational modification of the target that diminishes its interaction with an antibiotic is one of the classical ways how resistance develops^3, 38^. Yet, we found as remarkable that the modification of IleRS2 includes exchange of the most conserved 1^st^ and 3^rd^ positions^39^ of the class I AARS signature motif without compromising the variants housekeeping role and its broad distribution in bacteria (**Fig. 2**). In contrast, no IleRS1 with non-canonical signature motif has been found in nature so far (**Fig. 2**). Even more, introducing the non-canonical motif in laboratory-produced IleRS1 diminishes its activity (**Supplementary Fig. 3**). So, the signature motif exchange appears as an extraordinary resistance mechanism adopted exclusively in IleRS2. The insights into why IleRS2, but not IleRS1, can tolerate the non-canonical signature motif came from the crystal structures of the pair of PmIleRS1 and PmIleRS2, both in complex with Ile-AMS. Specifically, in PmIleRS2, both wild-type and the GXHH-mutant, the tip of the helix α1 carrying the HXGH or GXHH motif is rearranged (shifted) and the α-carbon of the third motif residue is displaced (away from the adenosine) by 1.7 Å relative to the analogous position in PmIleRS1 (**Fig. 3C**). This motif shift in IleRS2 solves the key problem associated with the change of a sequence, i.e., it enables accommodation of a larger amino acid, namely a His instead of the highly conserved Gly^39^, in the third position without a clash with the adenine base (**Fig. 4B**). In PmIleRS1, where no rearrangement of either HXGH or GXHH motif has been observed **(Fig. 3C, right)**, the histidine at the third position promotes mispositioning of the adenine moiety (**Fig. 4A**), resulting in the abolished activity of mut-GVHH-PmIleRS1 (**Supplementary Fig. 3**). That the steric constraints introduced by the 3^rd^ His promote a non-productive binding of ATP, which in turn diminishes IleRS1 activity, is further supported by mutational of analysis of the HLGH motif in methionyl-tRNA synthetase wherein Gly to Pro substitution induces severe *k*cat, but not *K*M (ATP), effect in the activation step and a loss of coupling binding energy between the amino acid substrate and ATP^40^.

Both *ileS1* and *ileS2* genes are proposed to be of ancient bacterial origin^17^ and have presumably been under selective pressure to develop resistance against naturally occurring mupirocin produced by *Pseudomonas fluorescens*^13^. However, did the same selective forces hold for both IleRS types? IleRS1 is found preferentially in faster- while IleRS2 in slower-growing bacteria^21^, raising the question whether a negative selection against catalytic trade-offs more prominently shaped IleRS1. That said, clinical isolates of *S. aureus*, including MRSA strains^34^, could evolve only a low resistance by IleRS1 mutations, while bacterial isolates with a higher IleRS1 resistance can be obtained exclusively in the laboratory because of their compromised fitness. Furthermore, some of the laboratory-selected *S. aureus* strains carry, among other mutations, also a His to Gln mutation at the 4^th^ position of the SaIleRS1 HIGH motif. Likely due to reduced catalytic properties, such mutations have not been observed in clinical isolates, explaining in part why, in spite of our considerable effort to engineer functional IleRS1 with a non-canonical motif, no active enzyme was obtained. Taken together, our results support a view that the differences between IleRS1 and IleRS2 are rooted deeply in the overall architecture of the catalytic domain, which is indicative for separate evolution trajectories of the IleRS1 and IleRS2 catalytic folds, likely because of distinct selection forces.

## Methods

### Cloning and mutagenesis

Preparation of the expression vectors encoding wild-type HIGH-EcIleRS1, HMGH-PmIleRS1 and HVGH-PmIleRS2 has been described elsewhere.^21, 41^. Expression vectors for wild-type ALHH- DrIleRS2 and HVGH-TtIleRS2 were prepared by inserting the coding sequences, PCR-amplified from genomic DNA of *Deinococcus radiodurans* R1 ATCC 13939 (DrIleRS2) and the plasmid TEx18A07 from the Riken BRC DNA Bank (TtIleRS2), into pET28b(+). The final constructs, which encoded non-codon optimized full-length enzyme coding sequences (CDS) fused to an N-terminal hexa-histidine tag, were verified by sequencing. Single point mutations were introduced by quick- change site directed mutagenesis (Q5-SDM kit (E0554, NEB)). Mutagenic primers were designed by the NEBaseChanger 1.3.3. software package that is available online. When suboptimal codon usage prohibited the mutagenic primer design (e.g. ALHH-DrIleRS2), cassette mutagenesis^42^ was used, where the parts of the original CDS were exchanged with synthetic cassettes carrying desired mutations (Twist Bioscience). The mutations were verified by sequencing.

### Protein expression and purification

Expression vectors carrying wild-type and mutant IleRSs were either transformed into chemically competent BL21(DE3) *E. coli* cells or, in the case of TtIleRS2-carrying plasmids, into Rosetta2 (DE3). For large-scale protein expression, cells were grown to early-log phase (0.4<OD<0.6) in LB broth supplemented with 1 mM MgCl2, which greatly improved the overall protein yield. Protein expression was induced by 0.25 mM IPTG (EcIleRS1, PmIleRS1, PmIleRS2, DrIleRS2 and their mutants) or 1 mM IPTG (wt- and mut-TtIleRS2) for 3h at 37 °C (wt and mut-EcIleRS), 5 h at 30 °C (wt and mut- PmIleRS1), or 16 h at 15 °C (DrIleRS2, TtIleRS2, PmIleRS2 and their mutants), followed by harvesting the cells using centrifugation and freezing the pellet at -80 °C. The latter significantly facilitated cell lysis efficiency and protein recovery.

The cell pellet from 0.5 L of culture was thawed at RT and resuspended in 10 ml IMAC A buffer (25 mM Hepes-KOH pH=7.5 at 20 °C, 500 mM NaCl, 10 mM imidazole, 10 mM 2- mercaptoethanol) followed by addition of 10 µg/ml DNase I, 10 µg/ml RNase I, 25 µg/ml lysozyme and 0.1 mM PMSF (final concentrations). The suspension was lysed by sonication (10 times for 45 seconds on 50 % sonication power with 1-min intervals between the pulses). The lysate was cleared by centrifugation (1 h at 25000 x g at 4 °C), and the supernatant containing soluble proteins was subjected to IMAC purification on a HisTrapHP 5 mL column (AktaPure 25 system). The column was washed with 20 CV of IMAC A and 10 CV IMAC B (25 mM Hepes-KOH pH=7.5 at 20 °C, 50 mM NaCl, 10 mM imidazole, 10 mM 2-mercaptoethanol) buffers, and the protein of interest was eluted by a linear gradient of imidazole using 20 CV of IMAC C buffer (25 mM Hepes-KOH pH=7.5 at 20 °C, 50 mM NaCl, 250 mM imidazole, 10 mM 2-mercaptoethanol). The fractions were pooled and the proteins further purified by IEX on a MonoQ 16/16 HR column equilibrated in IEX buffer A (25 mM Hepes-KOH pH=7.5 at 20 °C, 50 mM NaCl, 10 mM 2-mercaptoethanol). The unbound proteins were washed away with 20 CV IEX A, and elution of the protein of interest was achieved by a linear gradient of ionic strength using 20 CV IEX buffer B (25 mM Hepes-KOH pH=7.5 at 20 °C, 1000 mM NaCl, 10 mM 2-mercaptoethanol). The fractions containing pure protein were pooled, and the buffer was exchanged to storage buffer (25 mM Hepes-KOH pH=7.5 at 20 °C, 50 mM NaCl, 10 mM 2- mercaptoethanol). The proteins were concentrated to 16.5 mg/ml (HVGH-PmIleRS2 and mutants thereof), 20 mg/ml (HMGH-PmIleRS1, HIGH-EcIleRS1 and mutants thereof) or 10 mg/ml (HVGH- TtIleRS2 and mutants thereof), divided into 50 µl aliquots and flash frozen by plunging into liquid nitrogen. The purity of the final preparations is >95% as estimated by SDS-PAGE analysis.

### Determination of kinetic parameters in the IleRS activation reaction

Inhibition constants (*K*i) for mupirocin towards L-Ile and ATP in the activation step of the aminoacylation reaction catalyzed by wild-type and mutant IleRSs were determined by the ATP-PPi exchange assay^21, 43^. The reaction started by addition of the enzyme and was conducted at 30 °C in 20 µl reaction mixtures containing 55 mM Hepes-NaOH pH=7.5 (255 mM in case of ALHH-DrIleRS2), 30 mM MgCl2, 1 mM NaPPi, 5 mM DTT, 0.1 mg/ml BSA, ^32^P-PPi (0.2 - 0.4 µCi/µmol) and varying amounts of the enzyme, L-Ile, ATP and mupirocin. L-Ile and ATP concentrations ranged from 1/10 *K*M – 10 *K*M values. When measuring the *K*I towards one of the substrates, the other was held in saturation (concentrations at least 10 *K*M). The aliquots of the reaction mixture (1 – 2.5 μl) were quenched in 2 volumes of 500 mM NaOAc pH=4.5, 0,1% SDS, and formed ^32^P-ATP was separated from the remaining ^32^P-PPi by thin-layer chromatography on TLC plates (Marchanay-Nagel Polygram CEL 300 PEI UV254) using 750 mM KH2PO4 pH=3.5, 4 M urea buffer. Visualization and quantification of the signals was performed using a Typhoon Phosphoimager 5 (General Electric) and accompanying ImageQuant™ TL 10.2 analysis software. The kinetic constants were determined using a global fit option of the competitive inhibition model of GraphPad Prism 6 software. All experiments were performed at least in triplicates. The *k*cat was determined by dividing the maximal velocity, *V*M, with the used enzyme concentration.

### Protein crystallization and crystal stabilization

Prior to crystallization, the frozen protein samples were thawed and supplemented with the appropriate concentration of mupirocin (PmIleRS2 and mutants: 10 mM final concentration, PmIleRS1 and mutants: 3 mM final concentration), or Ile-AMS (2.5 mM final concentration). The solutions were centrifuged for 10 min at 4 °C and 25000 x g to remove particulates. The supernatants were used for sitting drop vapor diffusion crystallization experiments in 24-well Cryschem S Plates (Hampton) containing 300 µl of well solution as precipitant. The drops were set up by mixing 1 µl of protein:ligand solution with 1 µl of the well-solution, and crystals were obtained by incubation for one week at 4 °C, except for wt-PmIleRS1:mupirocin, which crystallized at 19 °C. Single crystals of wt- PmIleRS1:Ile-AMS or mut-GMHH-PmIleRS1:Ile-AMS complexes grew in crystallization buffer containing 300 mM Li2SO4, 150 mM Na2SO4, and 15 – 25% PEG3350. Crystal clusters of the wt- PmIleRS1:mupirocin complex were obtained from a crystallization buffer with 400 – 900 mM LiCl and 10 – 20% PEG3350. wt-PmIleRS2:Ile-AMS, wt-PmIleRS2:mupirocin, mut-GVHH-PmIleRS2:Ile- AMS and W130Q PmIleRS2:Ile-AMS crystallized in a buffer containing 0.2 – 0.5 M ammonium tartrate and 10 – 20% PEG3350. For wt-HMGH-PmIleRS1:Ile-AMS and mut-GMHH-PmIleRS1:Ile- AMS crystals, 20% glycerol (final concentration) was used as cryo-protectant, while for wt-HMGH- PmIleRS1:mupirocin, wt-HVGH-PmIleRS2:Ile-AMS, wt-HVGH-PmIleRS2:mupirocin, mut-GVHH- PmIleRS2:Ile-AMS and W130Q-PmIleRS2:Ile-AMS, crystals were stabilized and cryo-protected using PEG400 at a final concentration of 20%. Prior to data collection, all crystals were mounted on nylon loops and flash-frozen in liquid nitrogen.

### X- ray data collection and processing, structure determination, model building and refinement

Diffraction data of cryo-protected crystals were collected at 100 K using a wavelength of 1 Å at Beamline X06SA (PXI) or X06DA (PXIII) at the Paul Scherrer Institut (PSI) Villigen (Switzerland) and integrated and scaled using the XDS package^44^.

Initial phases for the wt-PmIleRS1:Ile-AMS complex were obtained by molecular replacement using the main body of SaIleRS1:tRNA:mupirocin (PDB 1QU2) and the TtValRS editing domain (PDB 1WKA) as search models. The structures of the mut-GMHH-PmIleRS1:Ile-AMS and wt-HMGH- PmIleRS1:mupirocin complexes were determined by molecular replacement using our solved wt- HMGH-PmIleRS1:Ile-AMS structure. Molecular replacement phases for the wt-HVGH- PmIleRS2:mupirocin dataset were obtained using the wt-HVGH-TtIleRS2:mupirocin structure (PDB 1JZS) as search model, and the structure was completed utilizing the *Phenix.autobuild* module in Phenix^45^ and manual model building in Coot 0.9.8.4.^46^. Initial phases for wt-HVGH-PmIleRS2:Ile- AMS, mut-GVHH-PmIleRS2:Ile-AMS, W130Q-PmIleRS2:Ile-AMS were obtained by molecular replacement using our wt-HVGH-PmIleRS2:mupirocin coordinates as search model.

All structures were finalized using iterative rounds of manual model building with Coot followed by coordinate and individual B-factor refinement in Phenix. To interpret the poorly resolved parts of the C-terminal tRNA-binding domain of both wt-HMGH-PmIleRS1 and mut-GMHH- PmIleRS1, we first docked an AlphaFold2^47^ model of this domain as a rigid body, manually adjusted the connections to the neighboring domains and refined the completed model as described above. Refinement and validation statistics for the final models are listed in **Supplementary Table 2**.

Inspection of our 1.9 Å mupirocin-bound PmIleRS2 structure clearly indicated an inverted chirality in the monic acid moiety of mupirocin. To account for the discrepancy between our ligand structure and the publicly available coordinate and topology files of the mupirocin (MRC) ligand, the chirality was manually changed in the restraints.

### Protein sequence retrieval for phylogenetic analysis

Sequences of the IleRS domains were retrieved using Pfam HMM profile (PF00133) from a set of 738 representative archaeal and bacterial genomes^24^ that span the prokaryote tree of life using HMMsearch with the Pfam-defined gathering threshold^48^. Incomplete sequences, i.e., those lacking the catalytic motif, were removed. Next, to remove redundancy, the remaining sequences were further clustered at 70% identity with CD-HIT^49^. Additionally, four sequences of ValRS – two bacterial and two archaeal – were selected from the species set and were used as outgroup. Overall, 374 sequences were taken and aligned using Mafft with the *–linsi* option^50^. The alignment was then trimmed with trimAl to remove gap-rich positions *(--gappyout* option)^51^.

### Phylogenetic analysis and ancestral sequence reconstruction

The resulting alignment was then used to build the phylogenetic tree of the family with the FastTree software with Jones-Taylor-Thornton (JTT) evolutionary models and the following parameters: *- pseudo, -spr 4, -mlacc 2, -slownni*^52^. The resulting tree was rooted with the ValRS outgroup. Ancestral sequence reconstruction was performed using *codeml* from the PAML package^53^ with the empirical JTT model and default parameters. Resulting trees were plotted and annotated using the program package ITOL^54^. The resulting tree, annotated with ancestral nodes, is shown in **Supplementary Fig 4** and **Supplementary data 2**.

### Molecular dynamics simulations

Four starting structures for molecular-dynamics (MD) simulations were used. In addition to the crystal structures of IleRS2 complexed to Ile-AMS or mupirocin, two apo-IleRS2 structures were generated by deleting the ligands. The parameters for mupirocin and Ile-AMS were prepared *de novo* (**Supplementary Fig. 15**), whereas the IleRS2 enzyme was described using the Amber ff14SB force field^55^. All models were dissolved in an explicit TIP3P water model^56^ and placed in a truncated octahedron-shaped simulation box. The systems were minimized in several steps and equilibrated at a temperature of 27 °C for 5 ns, followed by the production runs at the same temperature. Each model was simulated for 360 ns, and each trajectory consisted of 180 000 structures. The SHAKE algorithm was applied^57^, and a 2-fs step was used for numerical integration. The temperature was maintained using Langevin dynamics, and the pressure was controlled by Berendsen barostat^58^. The cut-off value was set to 9 Å. Production runs were performed with the AMBER20 on GPU using the pmemd.CUDA engine^59^.

### Molecular dynamics data analysis

The analysis was performed using the cpptraj module^60^ within the AmberTools20 package. The evolution of the secondary structure elements during the simulation time for helix α2 was monitored. Elements of the secondary structure were assigned according to the DSSP classification^61^ based on the analysis of φ and ψ torsion angles and hydrogen bonds.

### Synthesis of 5’-*O*-[(L-isoleucyl)sulfamoyl]adenosine (Ile-AMS)

Sulfamoyl chloride and compounds **1**-**3** (**Supplementary Fig. 16**) were prepared according to^62, 63^ with some changes detailed in **Supplementary methods**). Reagents and solvents for the synthesis of the compounds were obtained from Sigma-Aldrich Corp. (Germany) and Bachem (Switzerland). Organic solvents were further purified and/or dried using standard methods. Thin layer chromatography (TLC) was performed on Fluka silica gel (60 F254) plates (0.25 mm). Visualization was achieved using UV light at 254 nm and ninhydrine. Column chromatography was performed on Merck silica gel 60 (size 70-230 mesh ASTM). The ATR FT-IR spectrum was recorded on a FT-IR Perkin-Elmer Spectrum Two device (4000 do 400 cm^−1^ region). Mass spectra (ESI-MS) were recorded on an Agilent 6410 MS instrument. ^1^H and ^13^C NMR spectra of all precursors were recorded on a Bruker AV-III HD Bruker spectrometer at 400 MHz (^1^H) and 100 MHz (^13^C). All NMR experiments were performed at 298 K. Chemical shifts were referenced with respect to tetramethylsilane (TMS).

### Data availability

The coordinates and structure factors were deposited in the Protein Data Bank (PDB) under following accession numbers: **8C8U** (wt-HVGH-BmIleRS2:mupirocin), **8C8V** (wt-HVGH-BmIleRS2:Ile-AMS), **8C8W** (mut-GVHH-BmIleRS2:Ile-AMS), **8C9D** (wt-HVGH-BmIleRS2:Ile-AMS), **8C9E** (wt-HMGH- BmIleRS1:Ile-AMS), **8C9F** (mut-GMHH-BmIleRS1:Ile-AMS) and **8C9G** (wt-HMGH-BmIleRS1:mupirocin). Source data supporting Fig 2 (the sequence alignments in Fasta format and the trees in Newick format) are given in the Supplementary Data files 1 to 4. Source data supporting kinetic analysis presented in Table 1 are given in the Supplementary Data file 5. Starting molecular structures, topologies, and parameter files for molecular dynamics simulations are deposited on a publicly available GitHub repository accessible through the following link: https://github.com/aleksandra-mar/DATA_SHARING/releases/latest. Due to their large size, molecular dynamics trajectories are not publicly deposited but will be stored indefinitely on a personal data source. They will be disclosed without any restrictions upon request to the corresponding author. Other relevant data sources are either contained in the main manuscript or provided in the Supplementary Information file.

## Acknowledgment

This work was supported by the Swiss Enlargement Contribution in the framework of the Croatian-Swiss Research Programme, Grant IZHRZ0_180567 and European Regional Development Fund (infrastructural project CIuK, grant number KK.01.1.1.02.0016). A.M. acknowledges the Zagreb University Computing Centre (SRCE) for granting computational resources on the ISABELLA cluster. We would like to thank Jeff Errington for careful reading of the manuscript and Dan S. Tawfik, Dragana Despotovic and Liam M. Longo for numerous fruitful discussions.

## Authors contribution

I.G.S. and A.B. conceptualized and planned the experiments. A.B. cloned and purified the enzymes, performed kinetic and structural analyses, and prepared all figures. M.L. solved the crystal structure of the wt-HVGH-PmIleRS2:mupirocin complex and supervised the crystallographic work of A.B. J.J. performed the phylogenetic analysis and ancestral sequence reconstruction. V.Z. cloned mut- GMHH-BmIleRS1 and kinetically characterized the mut-GVHH-PmIleRS2 enzyme. V.P.P. and Z.C. synthesized the Ile-AMS analog. A.M. performed molecular dynamics simulations. I.G.S., A.B., M.L. and N.B analyzed the data. I.G.S. and N.B. conceived and supervised the project. I.G.S. wrote the manuscript with the contribution from all authors.

