## Supplementary material for "Antibiotic hyper-resistance in a class I aminoacyl-tRNA synthetase with altered active site signature motif": Suplementary information

**Supplementary Data 1.** Phylogenetic tree based on IleRS Pfam domain alignment (newick)

**Supplementary Data 2.** IleRS Pfam domain alignment (fasta)

**Supplementary Data 3.** The IleRS tree generated by the ancestral inference (newick)

**Supplementary Data 4.** Alignment of the ancestral IleRS sequences (fasta)

**Supplementary Data 5.** Source kinetic data (xlsx)

#### Supplementary methods

**Synthesis of 2',3'-*O*-isopropylidene-5'-*O*-sulfamoyladenosine (compound 1, Supplementary Fig. 18)**

Commercially available 2',3'-*O*-isopropylideneadenosine (723.2 mg, 2.35 mmol) in DME (70 ml) was placed under an argon atmosphere at 0 °C. The solution was stirred and NaH (141.2 mg, 3.53 mmol, 60% suspension in mineral oil) was added next. After 30 min the solution of freshly prepared sufamoyl chloride (see below) in DME (22 ml) was added dropwise. The resulting mixture was stirred overnight at room temperature. The reaction was monitored by TLC (EtOAc / MeOH, 10:1). Anhydrous methanol (35 ml) was added to the reaction mixture and it was evaporated *in vacuo*. Compound **1** was isolated as crude foam and used without purification in the next step.

To synthesize sulfamoyl chloride, anhydrous formic acid (133 μl, 3.53 mmol) was added dropwise to commercially available chlorosulfonyl isocyanate (307 μl, 3.55 mmol) at 0 °C. The mixture was stirred vigorously and the evolution of gases was observed during that process. After the addition of formic acid, the resulting solid was left at room temperature until gas evolution ceased. The structure of the product was confirmed by IR spectroscopy. The NCO band, a typical band for the starting chlorosulfonyl isocyanate, is missing from the IR spectrum of the product.

**Synthesis of 2',3'-*O*-isopropylidene-5'-*O*-[*N*-(*tert*-butyloxycarbonyl-L-isoleucyl) sulfamoyl] adenosine (compound 2)**

Compound **1** (342.2 mg, 0.886 mmol) was dissolved in DMF (8 ml) and stirred at room temperature under argon atmosphere. Commercially available Boc-L-isoleucine *N*-hydroxysuccinimide ester (291 mg, 0.886 mmol) and DBU (317.7 μl, 2.13 mmol) were added next. The reaction mixture was stirred for 2 h under the same conditions and monitored by TLC ($CHCl_3$ / MeOH, 3:1). The solvent and DBU were evaporated *in vacuo* using co-distillation with dry toluene. The residue was first triturated with dry diethyl ether and then with a 1:1 combination of diethyl ether / acetone. It was then purified using column chromatography ($CHCl_3$ / MeOH, 3:1).

**Synthesis of 5'-*O*-[(L-isoleucyl)sulfamoyl]adenosine (Ile-AMS, compound 3)**

To compound **2** (78 mg, 0.13 mmol), a combination of TFA and water (2 ml, 5:1 *v/v*) was added, and the resulting mixture was stirred at room temperature for 4 h. The reaction was monitored by TLC ($CHCl_3$ / MeOH, 3:1). Diethyl ether and absolute ethanol were added, and the mixture was evaporated *in vacuo*. The crude product was triturated with diethyl ether, and the residual was dissolved in 0.1 M aqueous ammonium formate and evaporated *in vacuo* again. The resulting Ile-AMS product (compound **3**) was purified by column chromatography ($CHCl_3$ / MeOH, 3:1).

### Supplementary tables

**Supplementary Table 1.** Mupirocin inhibition constants for the selected IleRS2 enzymes and their motif-exchanged mutants measured in the amino acid activation assay using ATP as a variable substrate.

| Enzyme | $K_i$ (MUP)$_{ATP}$ / μM # |
|---|---|
| wt-ALHH-DrIleRS2 | 7400 ± 500 |
| mut-HVGH-DrIleRS2 | 8.1 ± 0.5 |
| wt-HVGH-TtIleRS2 | 0.20 ± 0.01 |
| mut-GVHH-TtIleRS2 | 35 ± 2 |
| wt-HMGH-PmIleRS1† | 0.00029 ± 0.00015 |
| wt-HVGH-PmIleR2† | 1.04 ± 0.04 |
| mut-GVHH-PmIleRS2 | 480 ± 10 |

### The values represent the average value ± SEM of at least three independent experiments

† Values were taken from[1]

**Supplementary Table 2.** Summary of data collection and refinement statistics of the determined structures. Values in parentheses are for the highest resolution shell.

| **Structure**<br>**Ligand**<br>**PDB code** | **wt-PmIleRS1**<br>**Ile-AMS**<br>**8C9E** | **wt-PmIleRS1**<br>**mupirocin**<br>**8C9G** | **mut-GMHH PmIleRS1**<br>**Ile-AMS**<br>**8C9F** | **wt-PmIleRS2**<br>**Ile-AMS**<br>**8C9D** | **wt-PmIleRS2**<br>**mupirocin**<br>**8C8U** | **mut-GVHH PmIleRS2**<br>**Ile-AMS**<br>**8C8W** | **W130Q-PmIleRS2**<br>**Ile-AMS**<br>**8C9D** |
|---|---|---|---|---|---|---|---|
| Data collection statistics | | | | | | | |
| Space group | $P3_1 2\ 1$ | $P1\ 2_1\ 1$ | $P3_1 2\ 1$ | $P4_1 2_1 2$ | $P2_1 2_1 2$ | $P4_3 2_1 2$ | $P\ 4_3 2_1 2$ |
| Unit cell | 127.1 x 127.1 x 163.2 Å$^3$<br>α=β=90°, γ=120° | 66.8 x 144.3 x 108.4 Å$^3$<br>α=γ=90°, β=95.6° | 127.0 x 127.0 x 163.4 Å$^3$<br>α=β=90°, γ=120° | 108.1 x 108.1 x 240.0 Å$^3$<br>α=β=γ=90° | 89.6 x 124.8 x 114.5 Å$^3$<br>α=β=γ=90° | 107.9 x 107.9 x 239.4 Å$^3$<br>α=β=γ=90° | 108.2 x 108.2 x 240.4 Å$^3$<br>α=β=γ=90° |
| Res. range (Å) | 48.8 – 2.9 | 60.0 – 2.8 | 48.8 – 3.1 | 49.3 – 2.2 | 48.2 – 1.9 | 48.3 – 2.3 | 49.3 – 2.3 |
| No. of reflections | 713744 (74146) | 177079 (17665) | 582567 (60230) | 927990 (73039) | 1348929 (133421) | 835655 (74879) | 1645223 (131464) |
| Unique reflections | 34322 (2499) | 49643 (3810) | 28199 (1927) | 72948 (5848) | 101302 (7452) | 64159 (4825) | 64242 (5093) |
| Completeness (%) | 92.86 (74.11) | 90.26 (75.64) | 90.83 (69.78) | 94.10 (81.54) | 90.60 (74.27) | 92.59 (76.38) | 93.27 (81.00) |
| $R_{merge}$† | 0.06695 (3.219) | 0.15 (1.288) | 0.1224 (3.731) | 0.07646 (1.32) | 0.13 (2.579) | 0.1209 (2.125) | 0.1511 (2.303) |
| ⟨I│σ(I)⟩ | 30.66 (0.98) | 6.95 (0.94) | 19.70 (1.00) | 26.51 (1.66) | 14.93 (1.01) | 16.65 (1.20) | 22.62 (1.38) |
| Redundancy | 20.8 (22.0) | 3.6 (3.6) | 20.7 (21.8) | 12.7 (10.4) | 13.3 (13.4) | 13.0 (12.4) | 25.6 (21.0) |
| $CC_{1/2}$ | 1 (0.605) | 0.993 (0.373) | 0.999 (0.586) | 1 (0.62) | 0.999 (0.452) | 0.999 (0.454) | 0.999 (0.531) |
| Wilson *B* (Å$^2$) | 110.51 | 57.17 | 110.31 | 42.13 | 30.49 | 46.79 | 43.62 |
| $R_{work}$‡ (%) | 0.2421 (0.3894) | 0.1993 (0.3054) | 0.2395 (0.3840) | 0.1957 (0.2882) | 0.1872 (0.3283) | 0.1986 (0.3261) | 0.2019 (0.2977) |
| $R_{free}$§ (%) | 0.2685 (0.4452) | 0.2572 (0.3714) | 0.2766 (0.4368) | 0.2291 (0.3504) | 0.2273 (0.3605) | 0.2314 (0.3612) | 0.2416 (0.4005) |
| Molecules / ASU | 1 | 2 | 1 | 1 | 1 | 1 | 1 |
| No. atoms macromol. | 7408 | 14593 | 7408 | 8357 | 8223 | 8343 | 8338 |
| No. atoms solvent | 0 | 26 | 0 | 376 | 764 | 260 | 271 |
| No. atoms ligands | 106 | 91 | 108 | 73 | 73 | 73 | 73 |
| **Data refinement statistics** | | | | | | | |
| RMSD bonds (Å) | 0.002 | 0.002 | 0.002 | 0.003 | 0.008 | 0.003 | 0.003 |
| RMSD angles (°) | 0.430 | 0.44 | 0.440 | 0.530 | 0.890 | 0.510 | 0.620 |
| Rotamer outliers (%) | 0.87 | 0.87 | 1.00 | 0.97 | 1.25 | 0.65 | 0.66 |
| Clashscore | 8.8 | 5.0 | 8.4 | 3.8 | 3.1 | 4.4 | 5.0 |
| TLS groups | 7 | 0 | 7 | 0 | 0 | 0 | 0 |
| **Average *B* (Å$^2$)** | | | | | | | |
| All | 155.16 | 64.61 | 153.23 | 57.54 | 40.53 | 64.50 | 58.53 |
| Macromolecule | 155.34 | 64.67 | 153.36 | 57.83 | 40.34 | 64.78 | 58.87 |
| Waters | 0.00 | 50.73 | 0.00 | 50.14 | 42.64 | 53.00 | 48.04 |
| Ligands | 142.57 | 58.23 | 144.20 | 62.74 | 39.28 | 72.63 | 58.89 |
| **Ramachandran plot analysis (%)** | | | | | | | |
| Favored | 97.50 | 98.59 | 96.95 | 99.03 | 98.05 | 99.03 | 98.93 |
| Allowed | 2.50 | 1.41 | 3.05 | 0.97 | 1.86 | 0.87 | 1.07 |
| Outliers | 0.00 | 0.00 | 0.00 | 0.00 | 0.10 | 0.10 | 0.00 |

† $R_{merge} = \sum_{hkl}\sum_i |I_i(hkl) - \langle I(hkl)\rangle| / \sum_{hkl}\sum_i I_i(hkl)$. where $I_i(hkl)$ is the intensity of the $i^{th}$ measurement of the reflection *hkl* and $\langle I(hkl)\rangle$ is the mean value of $I_i(hkl)$ for all *i*.

‡ $R_{work} = \sum_{hkl}||F_{obs}| - |F_{calc}|| / \sum_{hkl}|F_{obs}|$. where $|F_{obs}|$ is the observed structure factor and $|F_{calc}|$ is the calculated structure factor.

§ $R_{free}$ is the same as $R_{cryst}$ except that it is calculated with a subset (5%) of data that were excluded from the refinement calculations.

#### Supplementary figures

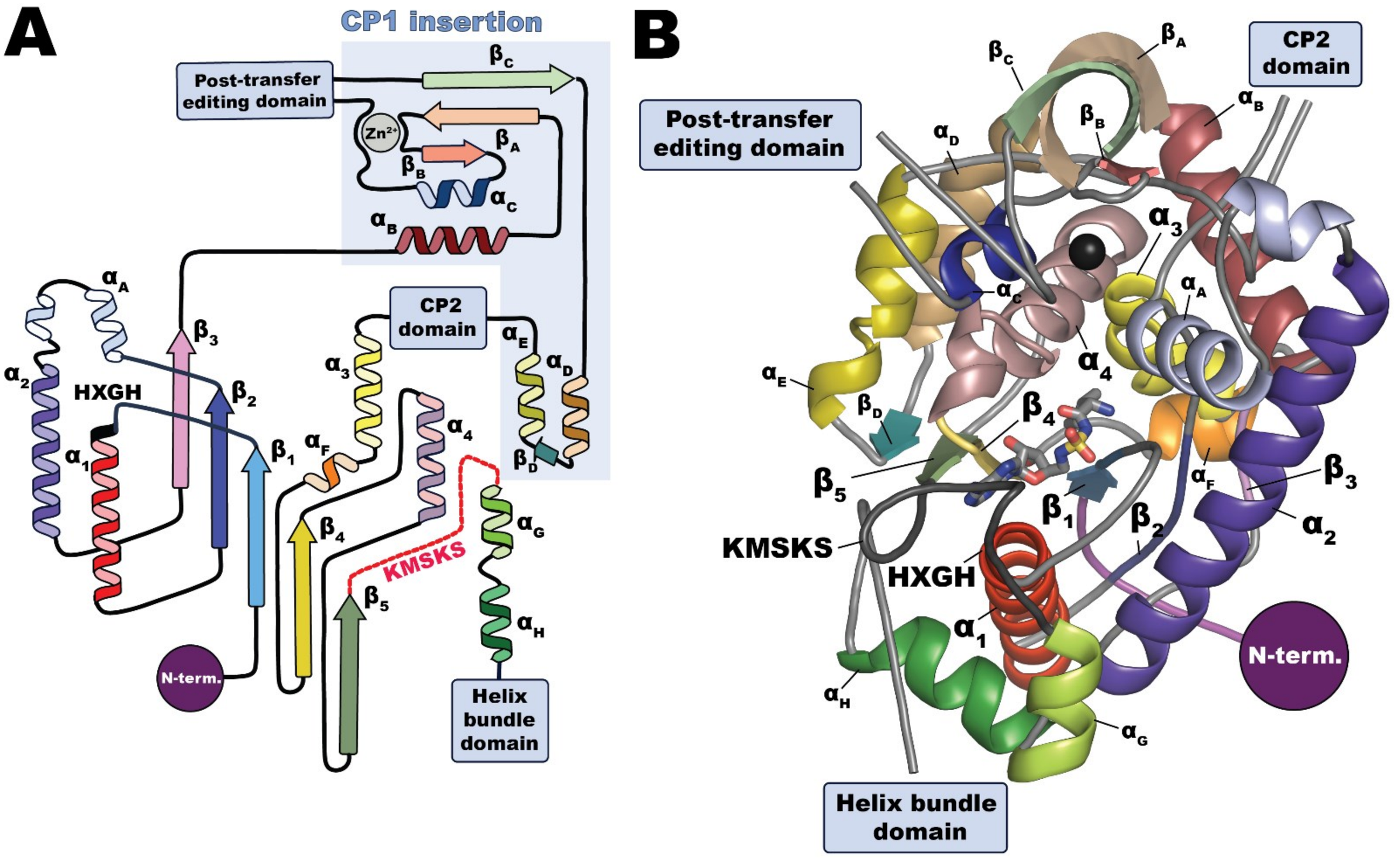

**Supplementary Fig. 1: Overview of the IleRS topology. (A)** The sandwich layer topology of the IleRS catalytic domain features five parallel β sheets bridged by four α helices. Catalytically important, class-defining signature motifs are located on the tip of the helix α1 (HXGH) and on the loop following the sheet β5 (KMSKS). **(B)** 3D representation of the TtIleRS2 (PDB code 1JZQ) catalytic domain bound to the analogue of isoleucyl-adenylate intermediate (Ile-AMS). The secondary structure elements are colored as in **(A)**. For clarity, the loops are shown in a simplified manner.

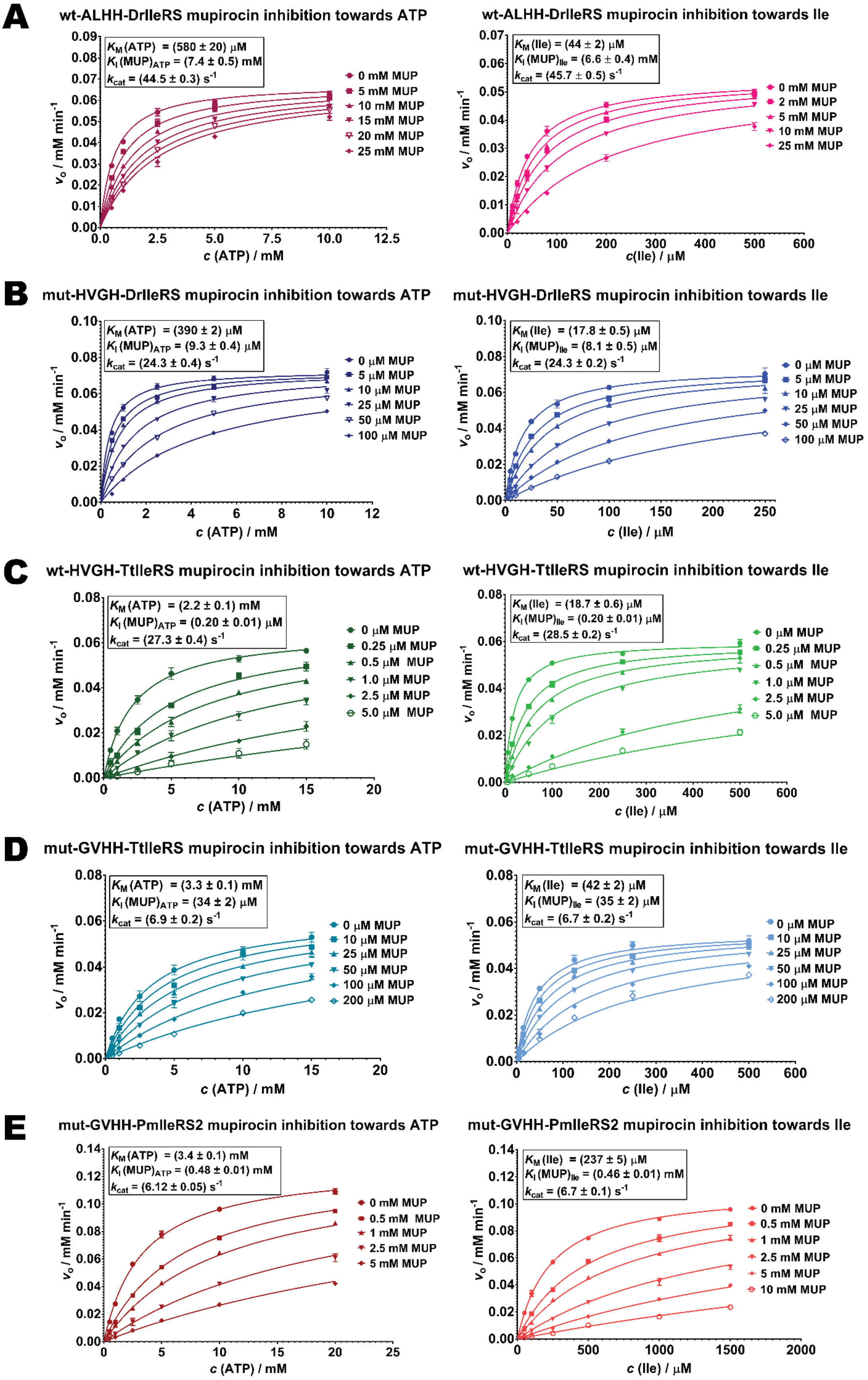

**Supplementary Fig. 2: Inhibition of IleRSs and their motif-exchanged mutants. (A-E)** Mupirocin (MUP) acts as a competitive inhibitor with highly similar inhibition constants against Ile and ATP in the amino acid activation step. The mupirocin concentration was varied in the range of 1 to 20 $K_I$, (1 to 5 $K_i$ in the case of wt-ALHH-DrIleRS), while the variable substrate (ATP or Ile) was varied in the range of 0.1 $K_M$ to 10 $K_M$. The other substrate (Ile or ATP) was kept in excess at 5 to 10 $K_M$. Data were globally fitted to the competitive inhibition model using GraphPad prism 6.01 for Windows software. The values represent the average value ± SEM of at least three independent experiments.

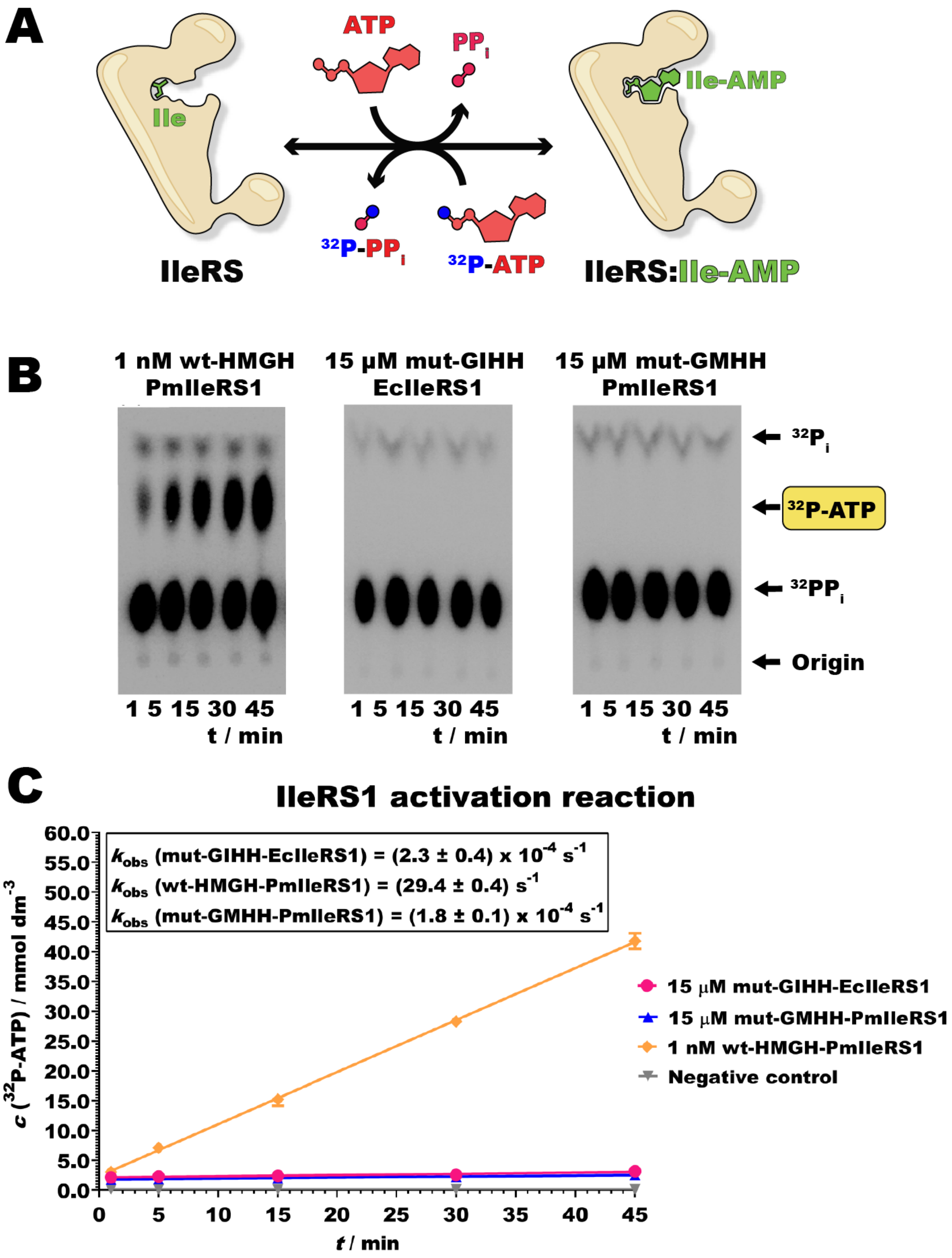

**Supplementary Fig. 3: Ile activation is abolished in mut-GMHH-PmIleRS1 and mut-GIHH-EcIleRS1. (A)** The activation is routinely measured by the ATP-PPi exchange assay[2]. **(B)** Thin-layer chromatograms showing the lack of $^{32}$P-ATP formation by the GXHH-IleRS1 enzymes present at high concentrations (15 µM) and prolonged reaction times. 1 nM wt PmIleRS1 was used as positive control. **(C)** Time courses of the data presented in (B). For the negative control, the enzymes were omitted from the reaction. The observed rate constant ($k_{obs}$) with wt-HMGH-PmIleRS1 agrees with the published data[1]. The $k_{cat}$ for wt-EcIleRS[3] is 56.7 ± 0.3 s$^{-1}$.

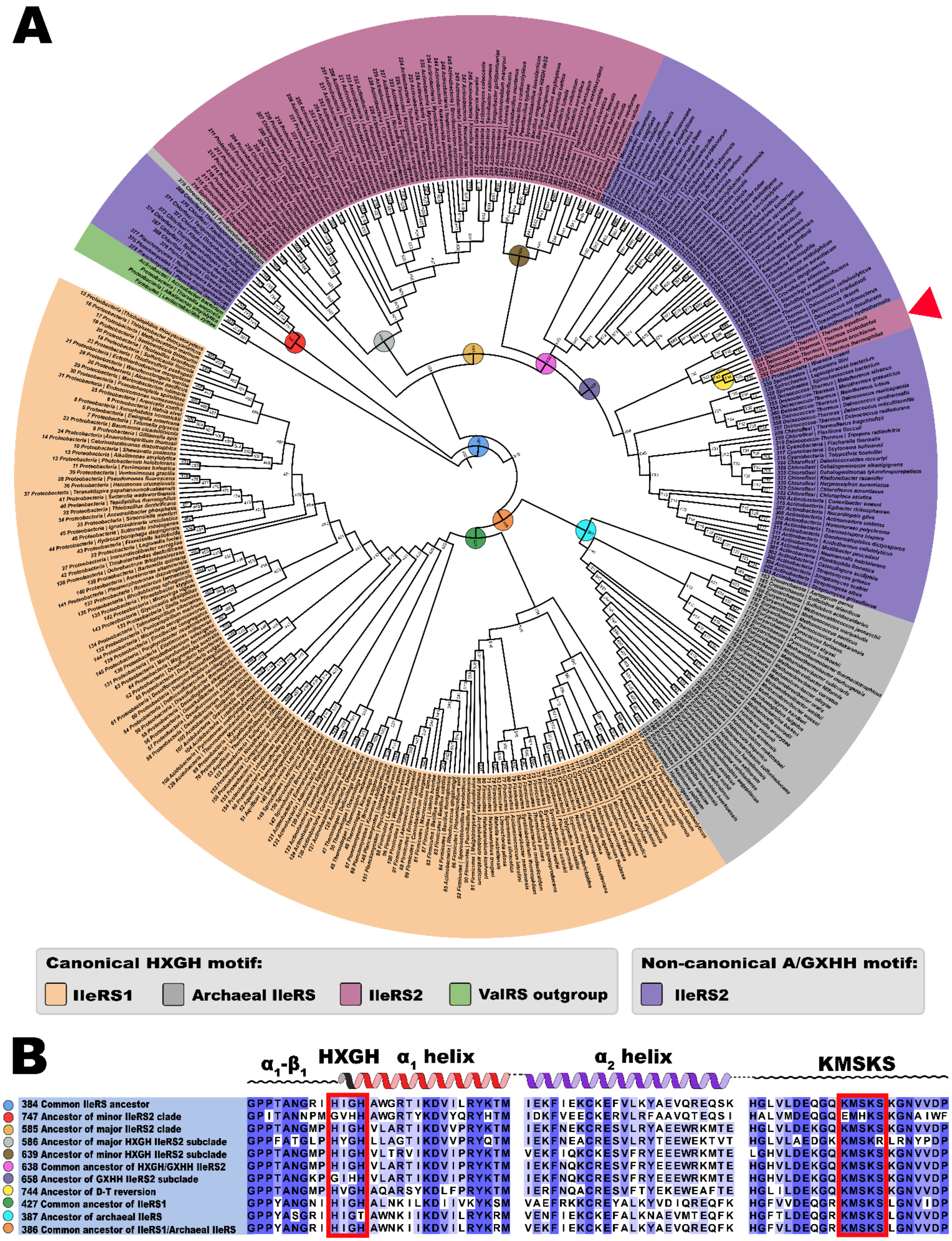

**Supplementary Fig. 4**: **Generation of an ancestral IleRS phylogenetic tree. (A)** An ancestral sequence reconstruction was performed by the PAML package[4] using a multiple sequence alignment of the PFAM domains of 379 IleRS sequences (158 HXGH IleRS1, 41 archeal HXG(H/T) IleRS, 77 HXGH IleRS2 and 86 GXHH IleRS2 sequences), and four ValRS sequences, used as an outgroup. The tree is plotted and annotated using the program package ITOL[5]. Naturally occurring reversion of the non-canonical to the canonical motif occurred in the *Deinococcus-Thermus* clade (red triangle). **(B)** Alignment of ancestral sequences of the selected nodes.

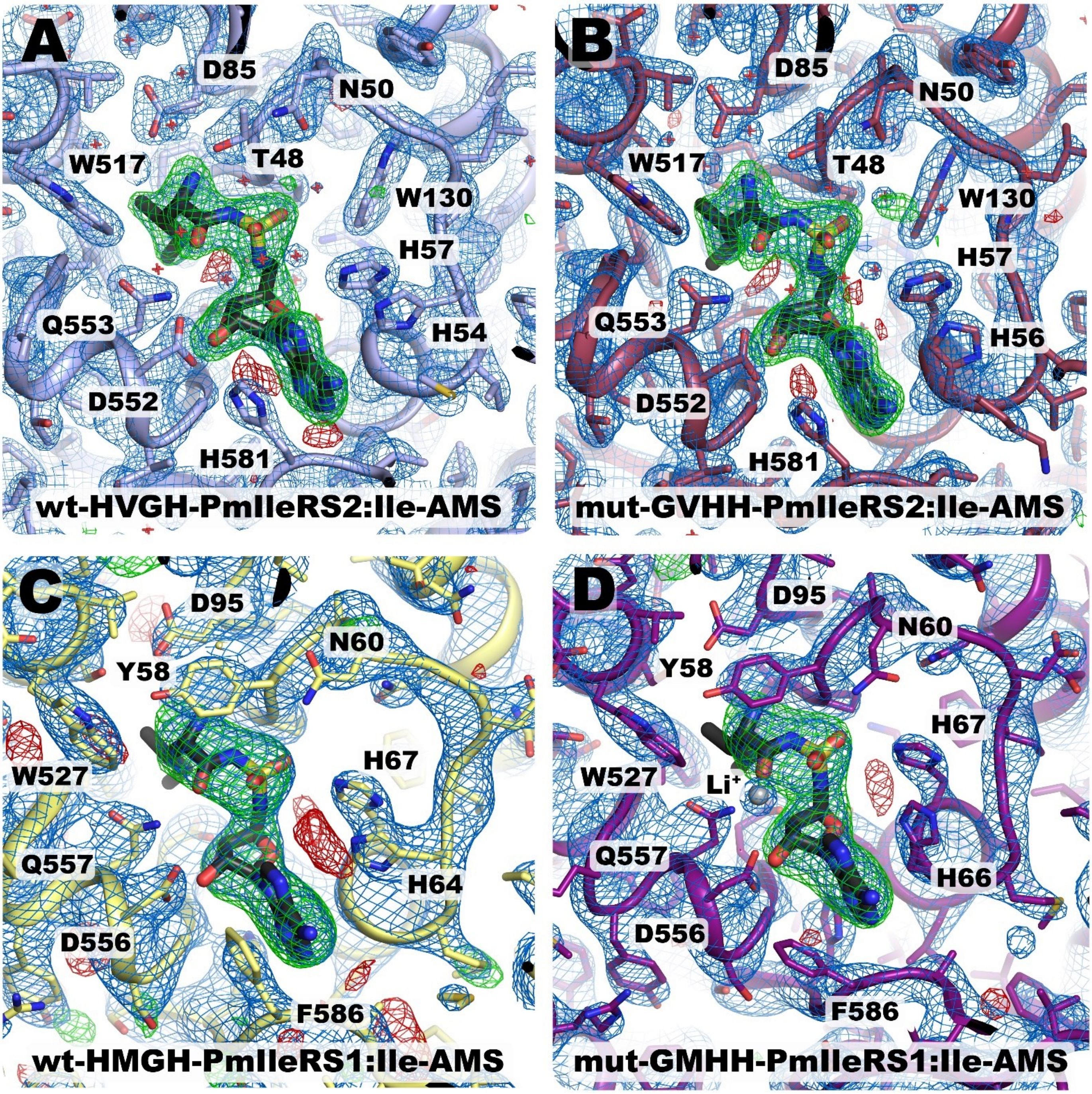

**Supplementary Fig. 5: Electron density and OMIT maps of Ile-AMS bound to PmIleRS1 and PmIleRS2. (A)** $2F_o$-$F_c$ Fourier difference map (blue) of the wt-HVGH-PmIleRS2:Ile-AMS complex. The map is resolved at 2.2 Å and contoured at a 2σ level. The unbiased $F_o$-$F_c$ Fourier difference OMIT map, contoured at 4σ (green) and -4σ (red) levels, shows the presence and conformation of the isoleucyl-adenylate analogue (Ile-AMS). **(B)** The $2F_o$-$F_c$ and corresponding $F_o$-$F_c$ OMIT Fourier difference density maps of the mut-GVHH-PmIleRS2:Ile-AMS unambiguously reveal the mutated signature motif residues and the bound Ile-AMS analogue. The map is resolved at 2.3 Å, with colors and contour levels as above. **(C)** 2.9 Å $2F_o$-$F_c$ Fourier difference density and corresponding $F_o$-$F_c$ OMIT maps of the wt-HMGH-PmIleRS1:Ile-AMS complex, contoured at 2σ and +/-2σ, respectively, and colored as above. **(D)** The $2F_o$-$F_c$ and corresponding $F_o$-$F_c$ OMIT Fourier difference density maps of the mut-GMHH-PmIleRS1:Ile-AMS unambiguously reveal the mutated signature motif residues and the bound Ile-AMS analogue. The map is resolved at 3.2 Å, with contour levels of 2σ for the $2F_o$-$F_c$ map and +/- 2σ levels for the respective $F_o$-$F_c$ OMIT maps. For generating the unbiased $F_o$-$F_c$ Fourier difference OMIT maps, the cofactors were removed from the models, followed by refinement until convergence (5 cycles) in Phenix after setting the B values to 100 Å$^3$ and randomizing the Cartesian coordinates by 0.1 Å.

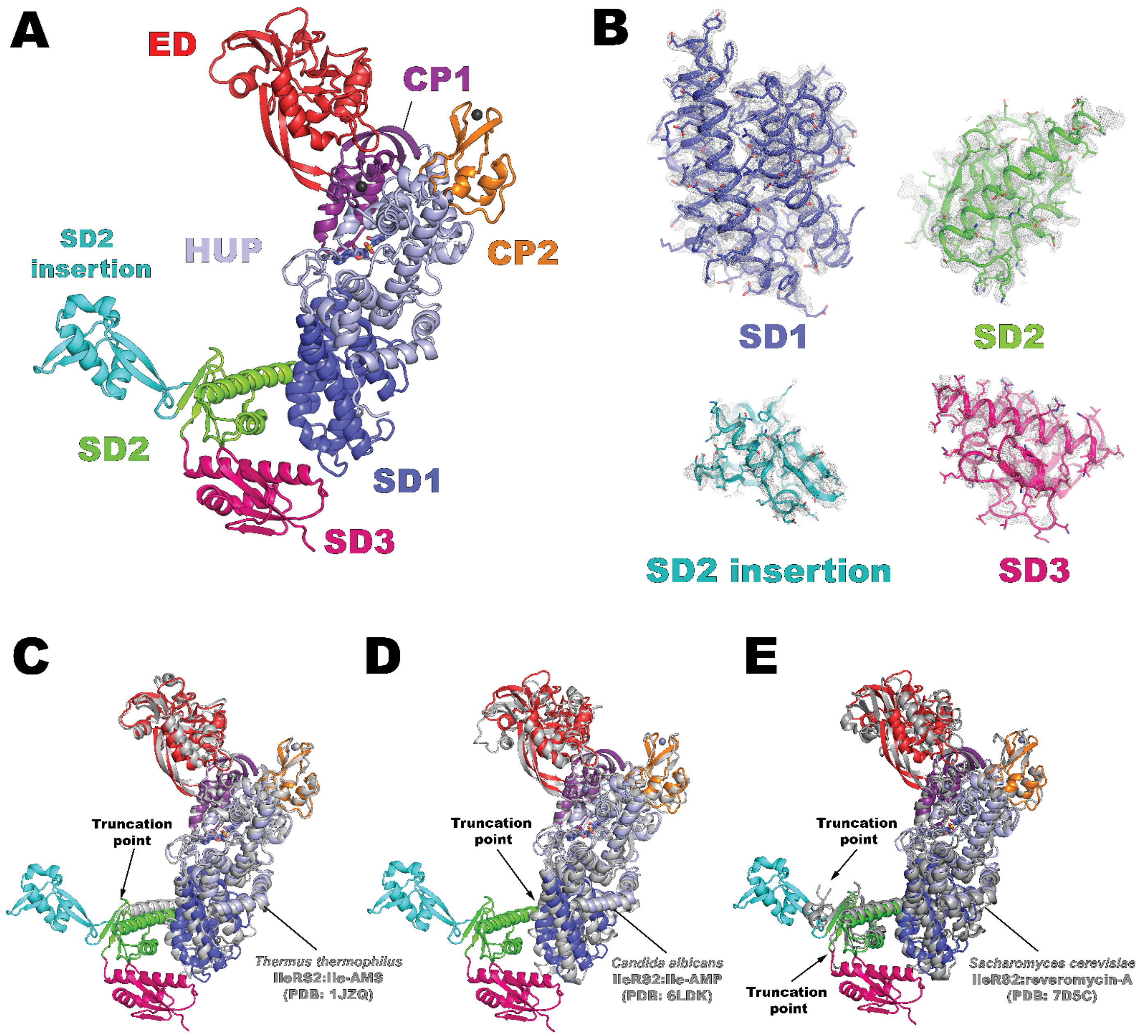

**Supplementary Fig. 6: Structure of the C-terminal domain of type 2 IleRS. (A)** In the wt-PmIleRS2:Ile-AMS structure, we were able to build for the first time the full-length C-terminal anticodon-binding domain of type 2 IleRS. **(B)** The domain consists of 3 subdomains. SD1 (helical bundle domain) and SD2 are structurally homologous to the corresponding subdomains in IleRS1. In IleRS2, the SD3 subdomain is homologous to SD2, while in IleRS1, the corresponding domain adopts a topologically unrelated αβ-fold with a zinc-binding motif (not shown). Further, IleRS2 SD2 features a novel insertion that adopts a βαβ-fold. The gray mesh around the models represents an unbiased composite OMIT map, contoured at a level of 2σ, showing the respective subdomains. **(C, D, E)** Comparison with the structures of *Thermus thermophilus* wt-IleRS2:Ile-AMS (PDB: 1JZQ), *Candida albicans* wt-IleRS2:Ile-AMP (PDB: 6LDK) and *Saccharomyces cerevisiae wt-IleRS2*:reveromycin (PDB:7D5C) depicts the novel parts of the structure unraveled by this study.

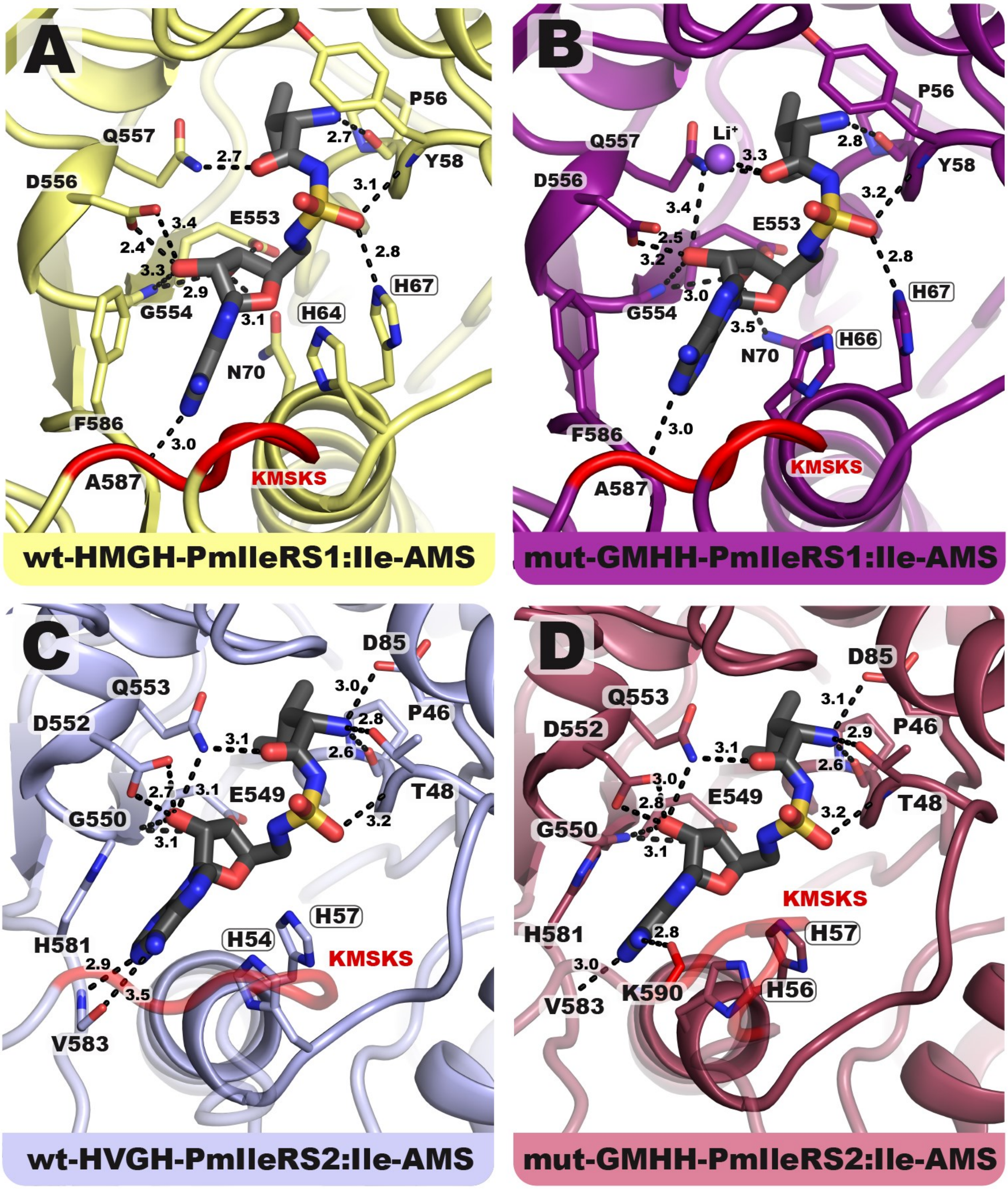

Supplementary Fig. 7: **Binding of Ile-AMS to wt-PmIleRS1, wt-PmIleRS2 and their signature motif mutants.** Ile-AMS binds to the WT enzymes **(A, C)** in an overall similar mode as to the corresponding mutants **(B, D)**. However, in the active site of the catalytically inactive mut-GMHH-PmIleRS1 **(B)**, the adenine base is mispositioned due to a steric constraint. See **Fig. 4** and main text for further details.

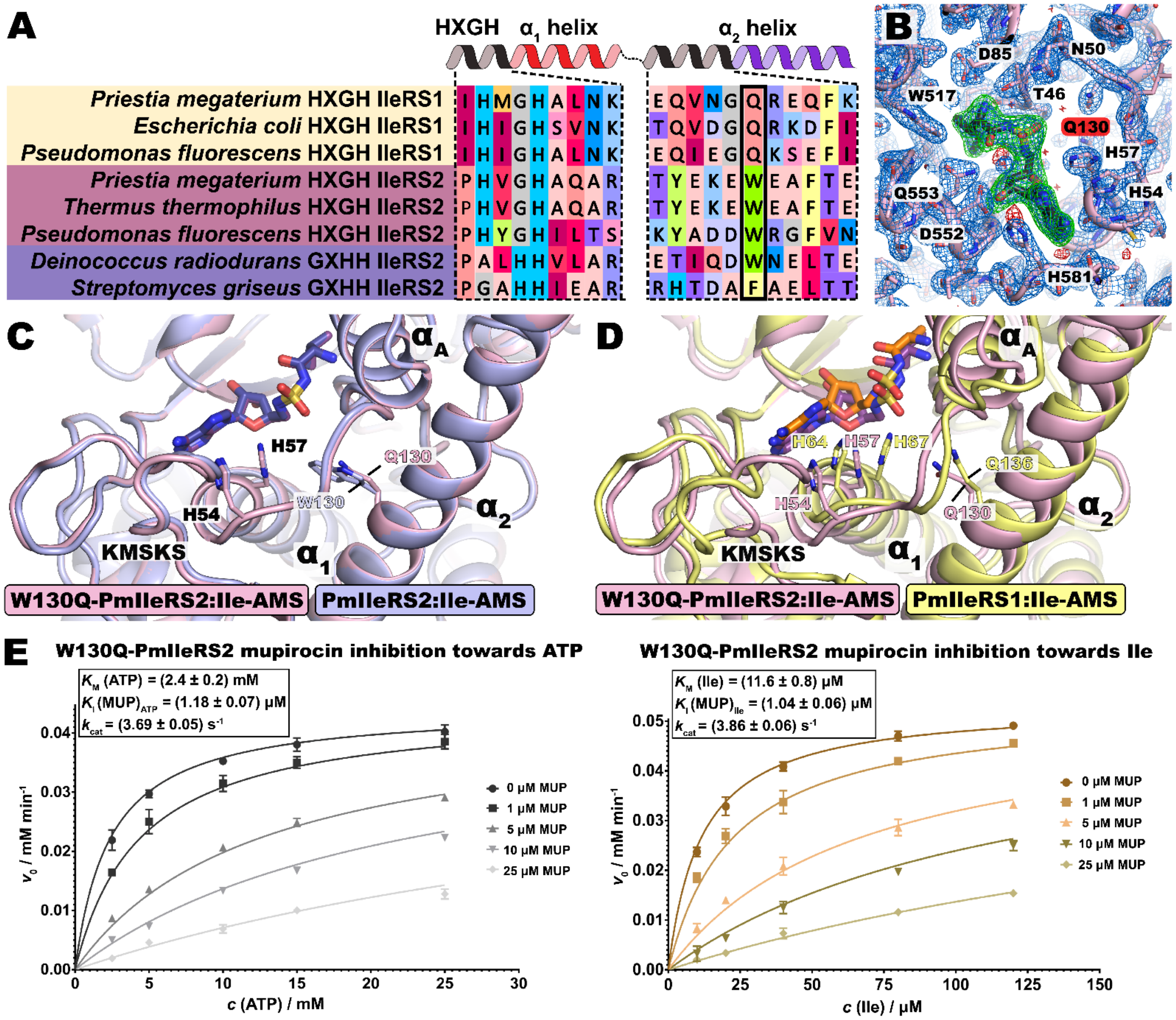

**Supplementary Fig. 8: The HXGH motif displacement is not dictated by a steric clash with a nearby α2 helix residue. (A, C, D, E)** Structural analysis identifies the α2 helix position (W130) that is in close proximity to the 4th His of the HXGH motif. A sequence alignment highlights this position as bulky in IRS2 (W130) but not in IRS1 (Q136), suggesting a crucial role of W130 in the displacement of the HXGH motif in IleRS2. However, in the structure of the W130Q PmIleRS2 mutant bound to Ile-AMS, the HXGH motif is at the same position of as in the WT enzyme (i.e. it remained replaced relative to IRS1:Ile-AMS). Kinetic analysis reveals that W130Q PmIleRS2 mutant displays an 18-fold lower $k_{cat}$ relative to the wild-type with a minor effect on the $K_M$ for Ile or ATP. The $K_i$ for mupirocin remained the same as for the wild-type enzyme. (**B**) A 2$F_o$-$F_c$ Fourier difference map of the W130Q-HVGH-PmIleRS2:Ile-AMS complex, resolved at 2.3 Å and contoured at a 2σ level, is shown in blue. The unbiased $F_o$-$F_c$ Fourier difference OMIT map is contoured at +/- 4σ (green/red), revealing the presence and conformation of the Ile-AMS analogue. For generating the unbiased $F_o$-$F_c$ Fourier difference OMIT map, the cofactor was removed from the model, followed by refinement until convergence (5 cycles) in Phenix after setting the B values to 100 Å$^3$ and randomizing the Cartesian coordinates by 0.1 Å.

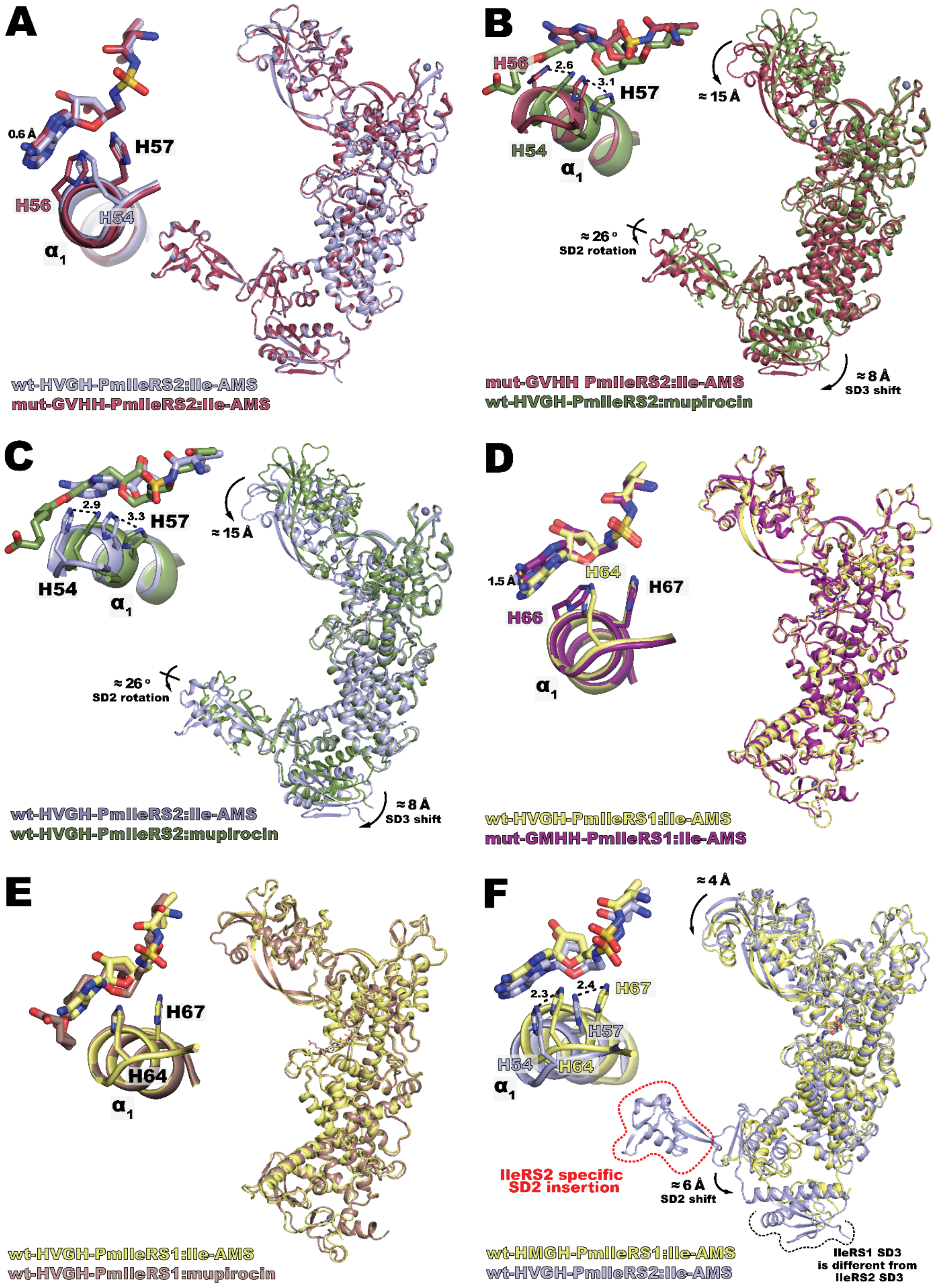

**Supplementary Fig. 9: Structural alignment of the wt-PmIleRS1, wt-PmIleRS2 and their signature motif mutants. (A-E)** Superposition of wt- and mut-PmIleRS1 or wt- and mut-PmIleRS2 full-length structures bound to Ile-AMS or mupirocin. The HXGH/GXHH motifs and bound ligands are highlighted. The cofactor-bound catalytic domains align well, revealing the conformational changes induced by binding of different ligands. A significant rearrangement of both the editing domain and C-terminal subdomains (SD2, SD3) relative to the catalytic domains are observed only for the pairs in B and C. **(F)** The superposition of wt-PmIleRS1 and wt-PmIleRS2 full-length structures shows that their catalytic and editing domains align well, while the C-terminal domains differ in topology and cannot be superimposed.

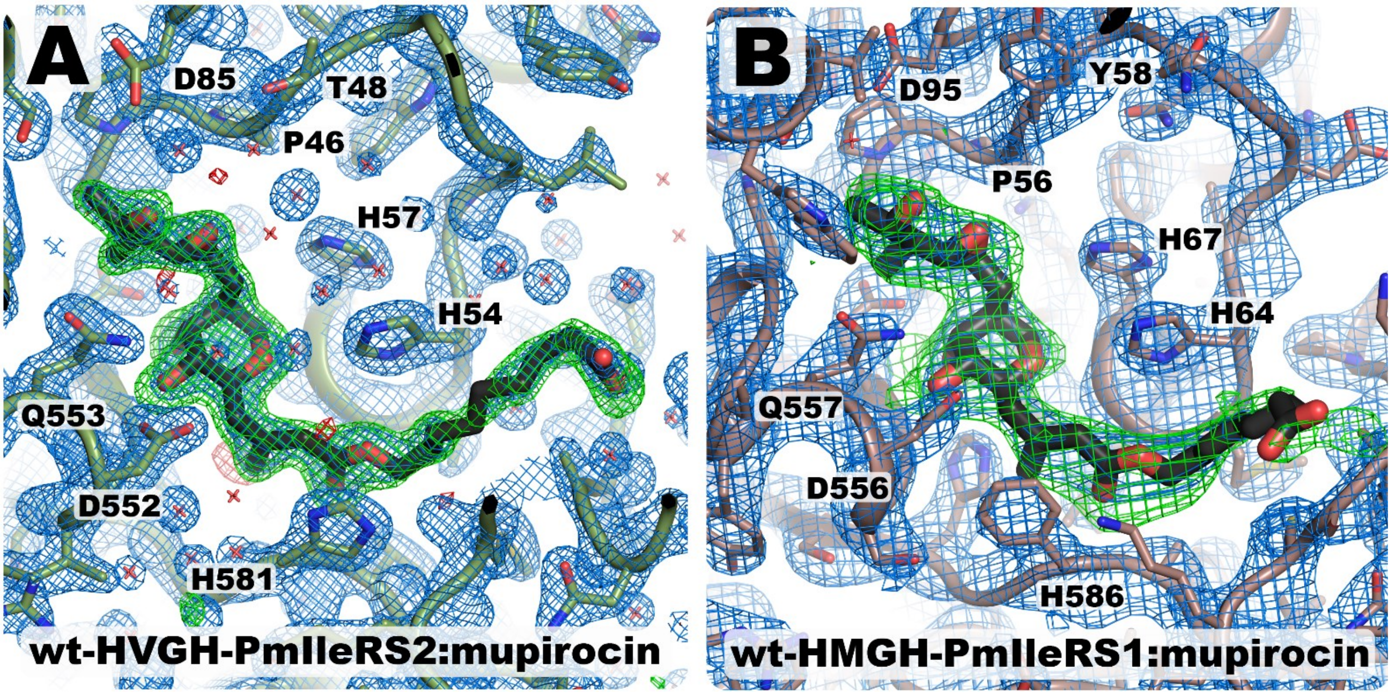

**Supplementary Fig. 10: Binding of mupirocin to wt-PmIleRS1 and wt-PmIleRS2. (A)** 2$F_o$-$F_c$ Fourier difference map (blue) of the wt-HVGH-PmIleRS2:mupirocin complex. The map is resolved at 1.9 Å and contoured at a 2σ level. The unbiased $F_o$-$F_c$ Fourier difference OMIT map, contoured at 5σ (green) and -5σ (red) levels, shows the presence and conformation of mupirocin. **(B)** A 2.95 Å 2$F_o$-$F_c$ Fourier difference density and corresponding $F_o$-$F_c$ OMIT maps of the wt-HMGH-PmIleRS1:mupirocin complex, contoured at 2σ and +/-2σ, respectively, are colored as above. For generating the unbiased $F_o$-$F_c$ Fourier difference OMIT maps, the cofactors were removed from the models, followed by refinement until convergence (5 cycles) in Phenix after setting the B values to 100 Å$^3$ and randomizing the Cartesian coordinates by 0.1 Å.

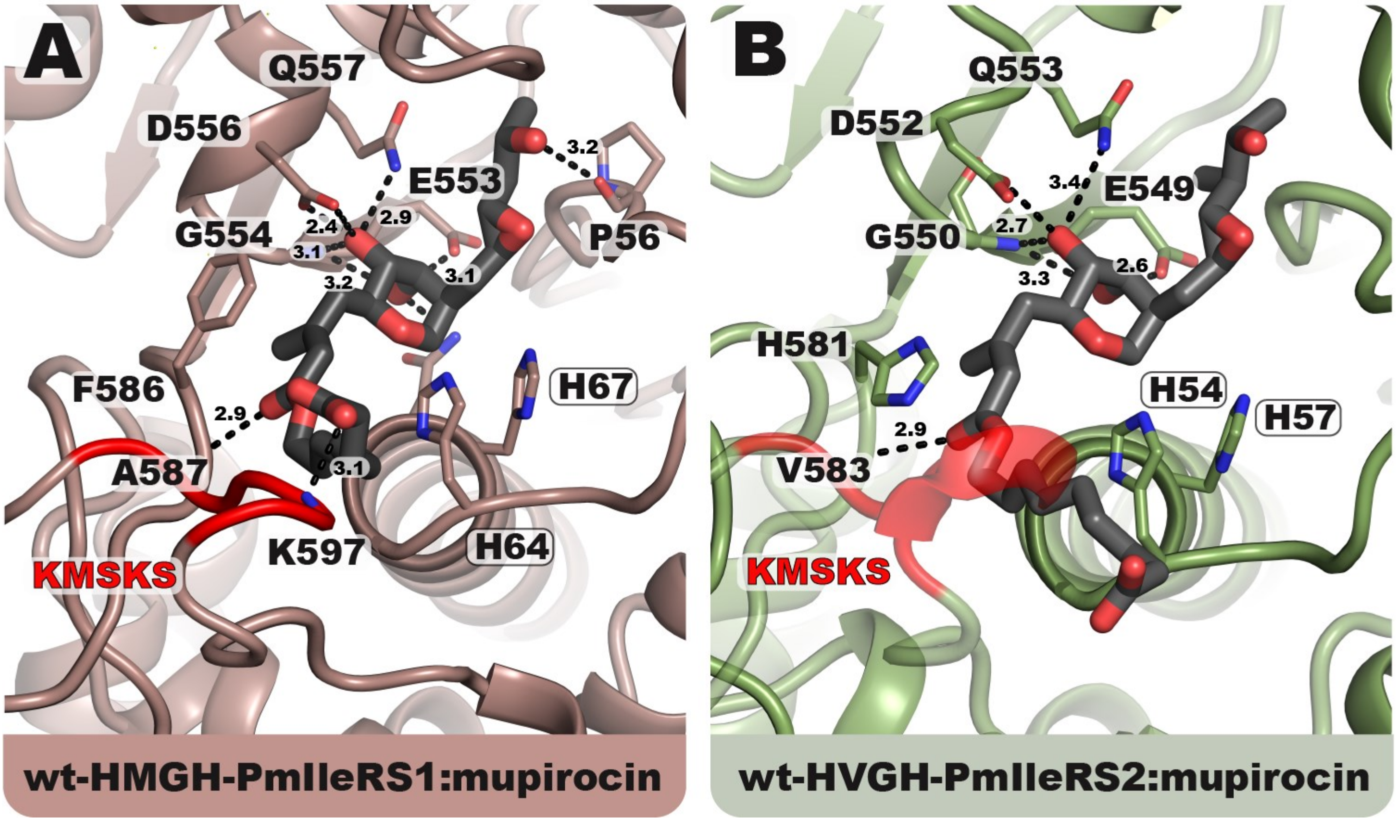

**Supplementary Fig. 11: Binding of mupirocin to wt-PmIleRS1 and wt-PmIleRS2. (A)** The monic acid portion of mupirocin binds to both enzymes in a canonical fashion with two additional interactions present in PmIleRS1, a hydrogen bond between Pro 56 and O13 and stacking of Phe 586 over the C2-C3 conjugated system. In PmIleRS1, the carboxylate of the nonanoic acid moiety of mupirocin establishes a hydrogen bond with NH backbone of Lys 597. **(B)** The closed conformation of the KMSKS loop in PmIleRS2 forces binding of the nonanoic acid moiety into the channel formed by the HXGH motif and KMSKS loop. This prevents the stabilizing interaction of the nonanoic carboxylate and Lys 597 from the KMSKS loop as observed in PmIleRS1.

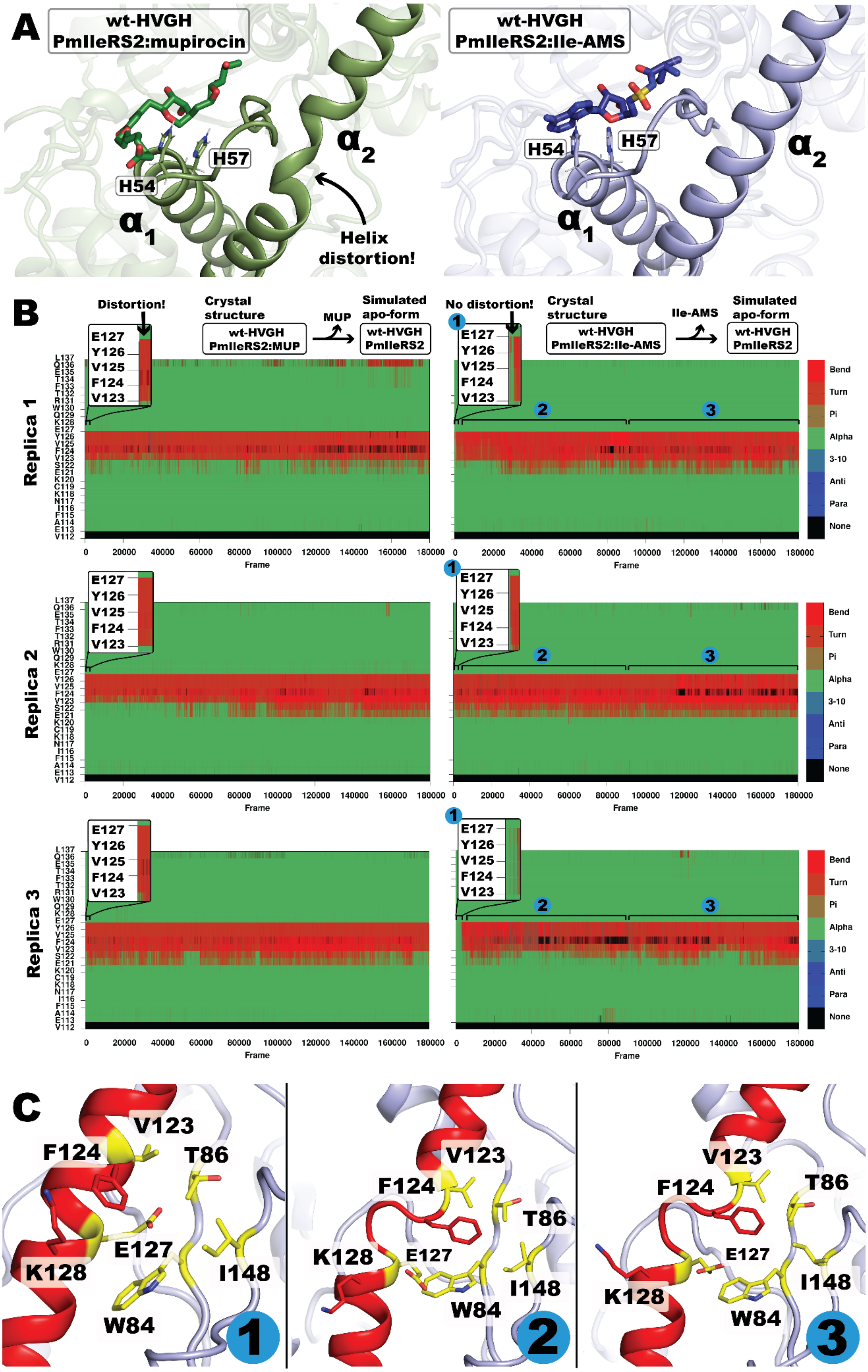

**Supplementary Fig. 12: MD simulations reveal the occurrence of a kink in helix α2 in PmIleRS2 independently of the presence of mupirocin or Ile-AMS. (A)** The comparison between crystal structures of wt-PmIleRS2 bound to mupirocin (left) or Ile-AMS (right) reveals a kink in the α2-helix only in the mupirocin-bound structure. **(B)** Secondary structures of helix α2 during triplicate 360 ns MD simulation time of wt-PmIleRS2 apo structures created by removing mupirocin from the wt-PmIleRS2:mupirocin crystal structure (left) or Ile-AMS from the wt-PmIleRS2:Ile-AMS crystal structure (right). The distortion in helix α2 originally present in the wt-PmIleRS2:mupirocin structure persists during the entire course of the MD simulation. In contrast, the undistorted helix α2 originally found in the wt-PmIleRS2:Ile-AMS crystal structure becomes distorted already after 5-10 ns of the simulation and remains distorted until its end. **(C)** Snapshots along the helix α2 distortion pathway. The distortion observed during the MD simulations might be driven by placement of residue Phe124 into the hydrophobic pocket formed by Ile148, Val 123, Trp 84, Thr 86 and Glu127, and by the loss of the cation-π interaction of Phe124 and Lys128 that is present in the wt-PmIleRS2:Ile-AMS structure. Taken together, the MD simulations suggest that the kink in helix α2 is an inherent feature of PmIleRS2 and is not related to mupirocin binding. It seems plausible to assume that two different conformations of the active site exist in solution and that binding of mupirocin or Ile-AMS pulls the equilibrium towards the conformations found in the crystal structures.

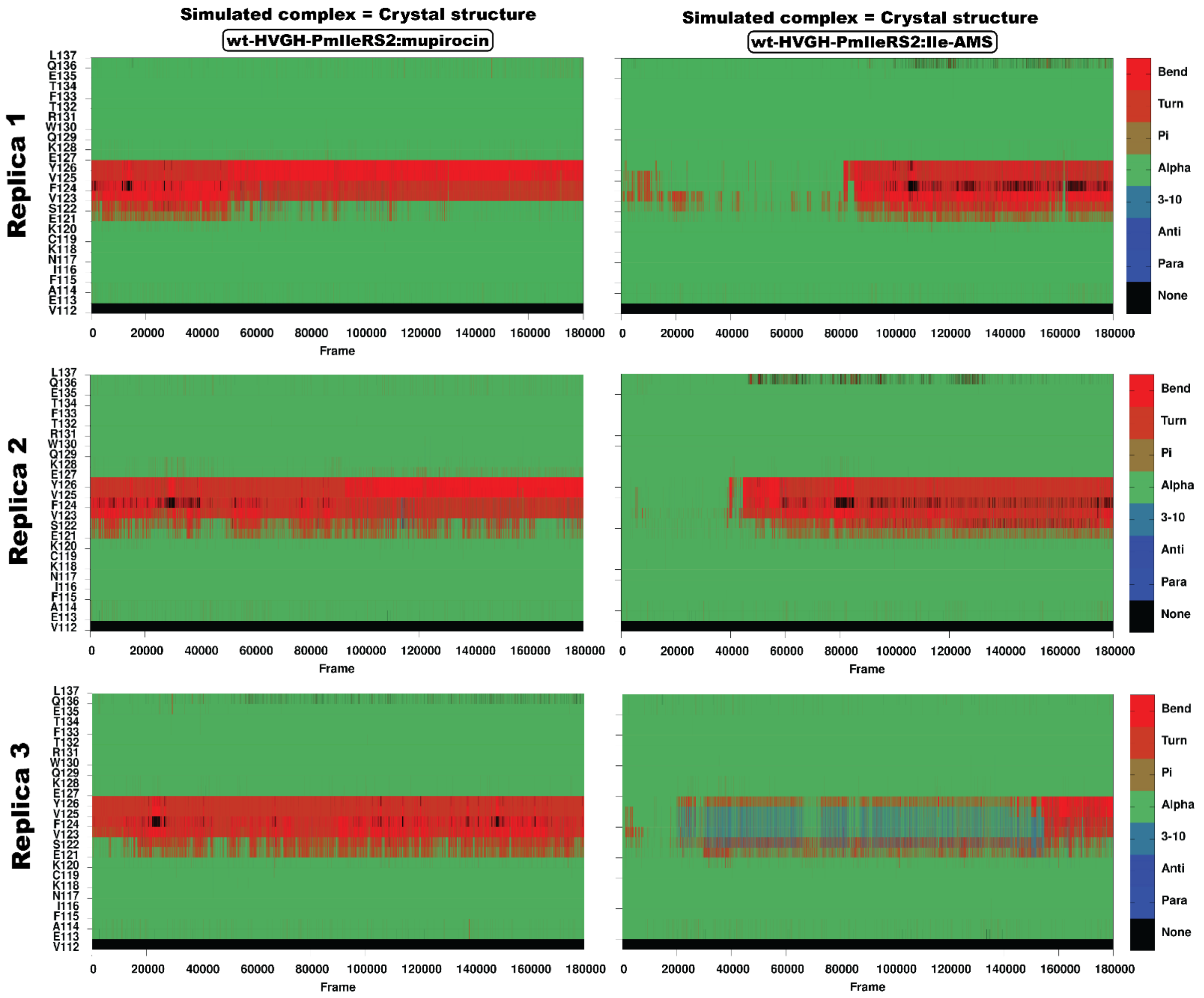

**Supplementary Fig. 13. Secondary structure monitoring of helix α2 during triplicate 360 ns MD simulations of the wt-PmIleRS2 bound to mupirocin (left) or Ile-AMS (right).** The distortion in helix α2 remains during the entire course of the simulation when mupirocin is bound. However, in the Ile-AMS-bound complex, the same distortion appears and remains stable for up to 300 ns. Ile-AMS remains bound to the enzyme even after helix α2 becomes distorted.

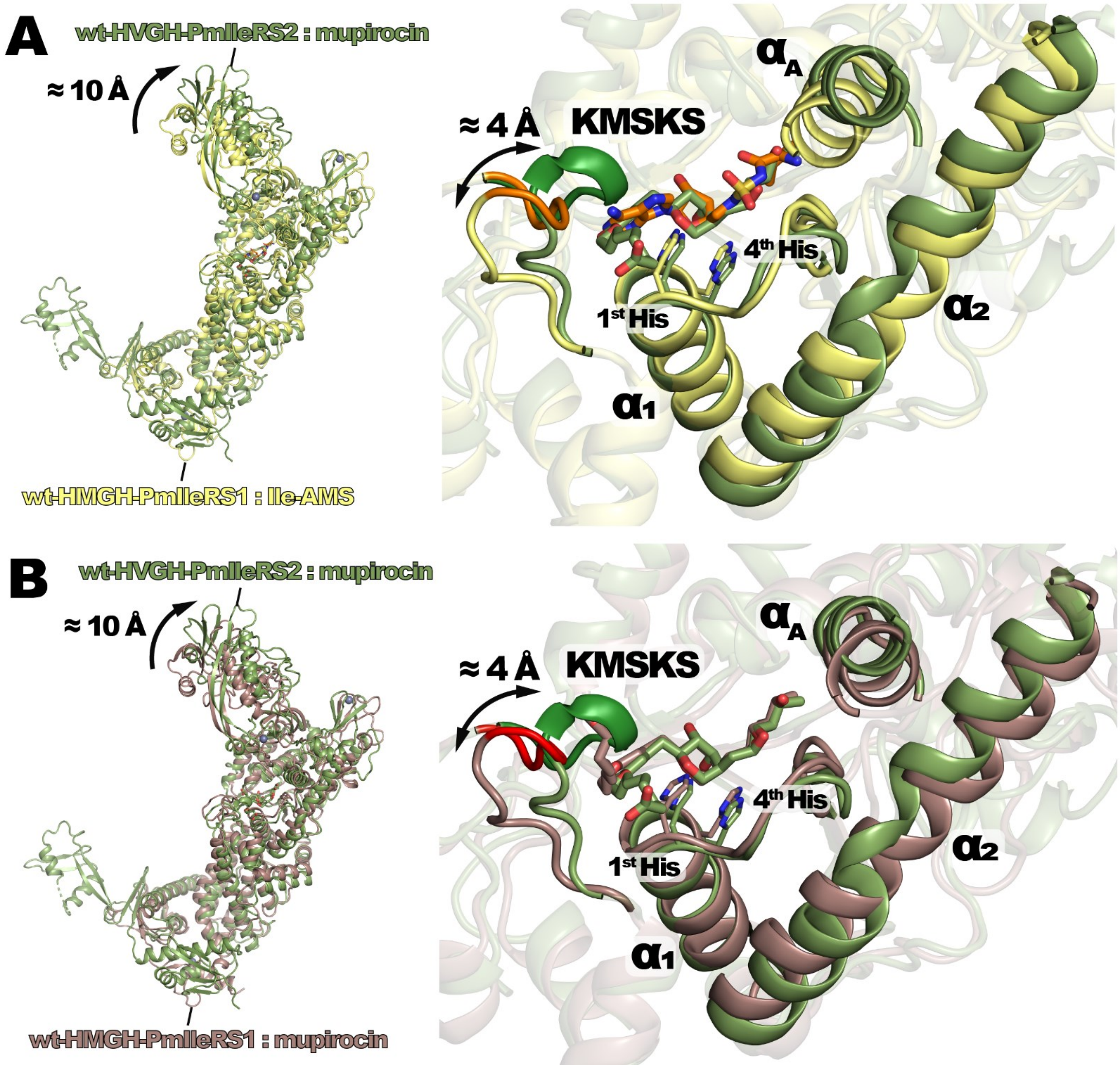

Supplementary Fig. 14: **Mupirocin binding to IleRS2 repositions the HXGH motif to the conformation observed in IleRS1.** The enzymes were superimposed using the backbone coordinates of the full-length structures (left panels) or the HUP catalytic cores (residues 51-174 and 521-635 in IleRS1 and 40-168 and 513-632 in IleRS2, right panels). **(A)** Structural overlay of PmIleRS1:Ile-AMS and PmIleRS2:mupirocin. The HXGH motif of the PmIleRS2:mupirocin complex assumes a similar conformation as the HXGH motif in PmIleRS1:Ile-AMS. Mupirocin binding to PmIleRS2 coincides with a distortion of helix α2, which is not present in PmIleRS1. Compared to PmIleRS1, the KMSKS loop of PmIleRS2 is around 4 Å closer to the active site. **(B)** Structural overlay of PmIleRS1:mupirocin with PmIleRS2:mupirocin. The overall superposition is very similar to the one presented in panel (A). In both cases, the catalytic cores align well (right panels), while whole enzymes superimpose poorly due to different orientations of the editing domain and the different topologies of their C-terminal domains (left panels). Helix α2 in the PmIleRS2:mupirocin structure is distorted compared to the one of PmIleRS1:mupirocin. In IleRS2, the closed arrangement of the KMSKS loop forces binding of the nonanoic moiety of mupirocin into a conformation that precludes formation of a hydrogen bond with Lys 597. See **Fig. 5** and main text for further details.

**A**

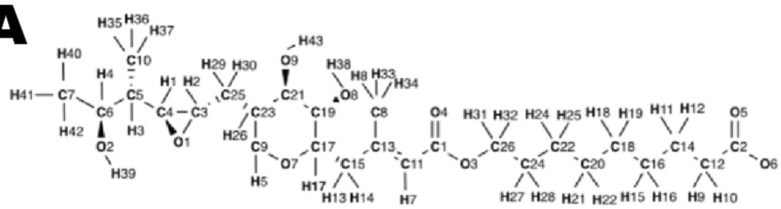

| Atom name | Atom type | Atomic charges |
|---|---|---|
| C7 | c3 | -0.192516 |
| H40 | hc | 0.045929 |
| H41 | hc | 0.045929 |
| H42 | hc | 0.045929 |
| C6 | c3 | 0.237954 |
| O2 | oh | -0.681418 |
| H39 | ho | 0.434490 |
| H4 | h1 | 0.088032 |
| C5 | c3 | 0.015898 |
| C10 | c3 | -0.27553 |
| H35 | hc | 0.077011 |
| H36 | hc | 0.077011 |
| H37 | hc | 0.077011 |
| H3 | hc | 0.066074 |
| C4 | cx | 0.037224 |
| O1 | os | -0.391945 |
| H2 | h1 | 0.100980 |
| C3 | cx | 0.028980 |
| H1 | h1 | 0.112393 |
| C25 | c3 | -0.142991 |
| H29 | hc | 0.070817 |
| H30 | hc | 0.070817 |
| C23 | c3 | 0.041905 |
| C21 | c3 | 0.174318 |
| C19 | c3 | 0.036021 |
| O8 | oh | -0.702779 |
| H38 | ho | 0.466052 |
| H20 | h1 | 0.131083 |
| O9 | oh | -0.695301 |
| H43 | ho | 0.468355 |
| H23 | h1 | 0.087215 |
| H26 | hc | 0.089134 |
| C9 | c3 | -0.021985 |
| H5 | h1 | 0.088118 |
| H6 | h1 | 0.088118 |
| O7 | os | -0.414575 |
| C17 | c3 | 0.101901 |
| H17 | h1 | 0.118525 |
| C15 | c3 | -0.06566 |

| Atom name | Atom type | Atomic charges |
|---|---|---|
| H13 | hc | 0.058559 |
| H14 | hc | 0.058559 |
| C13 | c2 | 0.020052 |
| C8 | c2 | -0.40299 |
| H33 | ha | 0.118493 |
| H34 | ha | 0.118493 |
| H8 | hc | 0.118493 |
| C11 | c3 | -0.054018 |
| H7 | hc | 0.043744 |
| C1 | c | 0.675721 |
| O4 | o | -0.584931 |
| O3 | os | -0.325571 |
| C26 | c3 | -0.088394 |
| H31 | h1 | 0.120627 |
| H32 | h1 | 0.120627 |
| C24 | c3 | -0.054859 |
| H27 | hc | 0.046484 |
| H28 | hc | 0.046484 |
| C22 | c3 | 0.018468 |
| H24 | hc | -0.009695 |
| H25 | hc | -0.009695 |
| C20 | c3 | 0.020990 |
| H21 | hc | -0.031662 |
| H22 | hc | -0.031662 |
| C18 | c3 | 0.000154 |
| H18 | hc | -0.000695 |
| H19 | hc | -0.000695 |
| C16 | c3 | 0.004906 |
| H15 | hc | -0.003002 |
| H16 | hc | -0.003002 |
| C14 | c3 | 0.007274 |
| H11 | hc | -0.001036 |
| H12 | hc | -0.001036 |
| C12 | c3 | -0.035303 |
| H9 | hc | -0.037749 |
| H10 | hc | -0.037749 |
| C2 | c | 0.838468 |
| O6 | o | -0.830688 |
| O5 | o | -0.830688 |

**B**

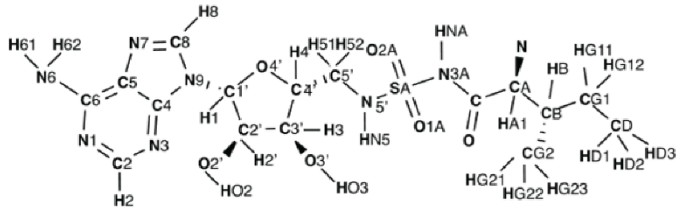

| Atom name | Atom type | Atomic charges |
|---|---|---|
| N | n3 | -0.916800 |
| CA | c3 | 0.121500 |
| C | c | 0.651100 |
| O | o | -0.629100 |
| CB | c3 | -0.057700 |
| CG1 | c3 | -0.081400 |
| CG2 | c3 | -0.090100 |
| CD | c3 | -0.103100 |
| C1' | c3 | 0.255800 |
| C2 | ca | 0.688900 |
| C2' | c3 | 0.076100 |
| C3' | c3 | 0.085100 |
| C4 | ca | 0.475800 |
| C4' | c3 | 0.092100 |
| C5 | ca | -0.238200 |
| C5' | c3 | 0.221800 |
| C6 | ca | 0.763200 |
| C8 | cc | 0.389700 |
| N1 | nb | -0.794000 |
| N3 | nb | -0.732000 |
| N3A | n | -0.671300 |
| N5' | n3 | -0.915100 |
| N6 | nh | -0.957500 |
| N7 | nd | -0.530100 |
| N9 | na | -0.284000 |
| O1A | o | -0.633800 |
| O2' | oh | -0.587800 |
| O2A | o | -0.633800 |
| O3' | oh | -0.593800 |

| Atom name | Atom type | Atomic charges |
|---|---|---|
| O4' | os | -0.417600 |
| SA | s6 | 1.573500 |
| HG23 | hc | 0.044700 |
| HG22 | hc | 0.044700 |
| HG21 | hc | 0.044700 |
| HG12 | hc | 0.037700 |
| HG11 | hc | 0.037700 |
| HO3 | ho | 0.421000 |
| HO2 | ho | 0.422000 |
| HNA | hn | 0.332500 |
| HN5 | hn | 0.471800 |
| HN2 | hn | 0.369300 |
| HN1 | hn | 0.369300 |
| HD3 | hc | 0.043700 |
| HD2 | hc | 0.043700 |
| HD1 | hc | 0.043700 |
| HB | hc | 0.058700 |
| HA1 | h1 | 0.101700 |
| H8 | h5 | 0.094800 |
| H62 | hn | 0.448800 |
| H61 | hn | 0.448800 |
| H52 | h1 | 0.068700 |
| H51 | h1 | 0.068700 |
| H4 | h1 | 0.082700 |
| H3 | h1 | 0.098700 |
| H2' | h1 | 0.087700 |
| H2 | h5 | 0.052100 |
| H1 | h2 | 0.134700 |

**Supplementary Fig. 15. Atom names, atom types and atomic charges used for the ligands during MD simulations. (A)** Mupirocin **(B)** Ile-AMS, 5′-N-[N-(L-isoleucyl)sulfamoyl]adenosine

**A**

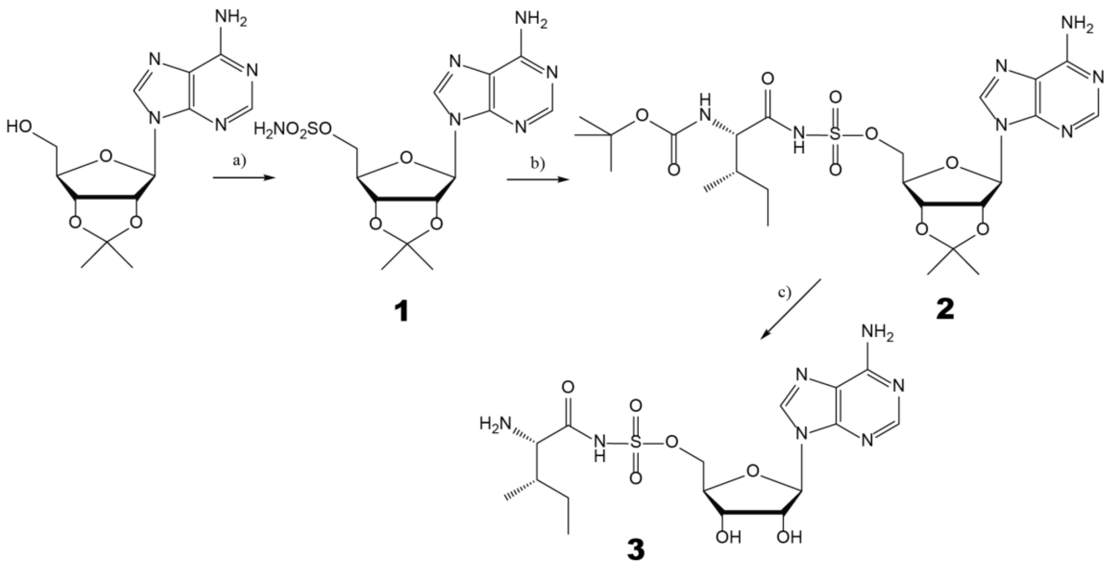

**B**

#### Compound 1 $^1$H NMR spectrum

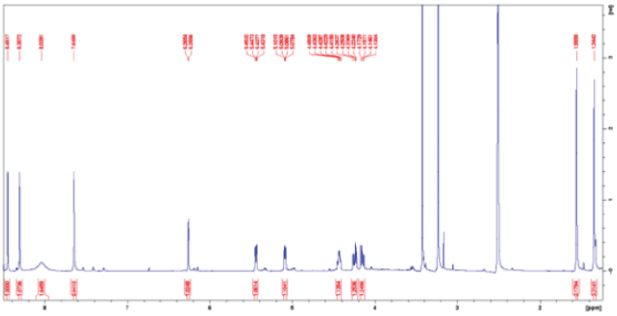

**$R_f$ = 0.41 (EtOAc / MeOH, 10 : 1). $^1$H NMR (DMSO-$d_6$) δ/ppm: 8.45 (s, 1H, H-8); 8.30 (s, 1H, H-2); 8.04 (br s, 2H, $SO_2NH_2$); 7.65 (s, 2H, $NH_2$); 6.26 (d, 1H, $J_{1',2'}$= 2.3 Hz, H-1'); 5.44 (dd, 1H, $J_{2',3'}$ = 6.3 Hz, $J_{2',1'}$ = 2.4 Hz, H-2'); 5.09 (dd, 1H, $J_{3',2'}$ = 6.2 Hz, $J_{3',4'}$ = 3.1 Hz, H-3'); 4.51-4.42 (m, 1H, H-4'); 4.24 (dd, 1H, $J_{5a',4'}$ = 5.2 Hz, $J_{5a',5b'}$ = 10.9 Hz, H-5a'); 4.15 (dd, 1H, $J_{5b',4'}$ = 6.3 Hz, $J_{5a',5b'}$ = 10.7 Hz, H-5b'); 1.56 (s, 3H, $CH_3$); 1.34 (s, 3H, $CH_3$). ESI-MS: *m*/*z* $[M+H]^+$ calc for $C_{13}H_{18}N_6O_6S$: 386.1, found: 387.3.**

#### Compound 2 $^1$H NMR spectrum

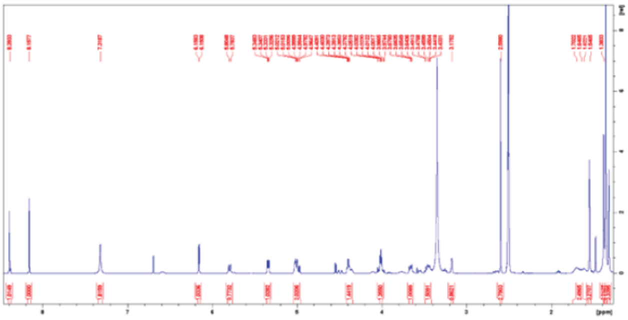

**Yield: 318 mg (60%). White crude foam. $R_f$ = 0.58 ($CHCl_3$ / MeOH, 3: 1). $^1$H NMR (DMSO-$d_6$) δ/ppm: 8.39 (s, 1H, H-8); 8.16 (s, 1H, H-2); 7.32 (br s, 2H, $NH_2$); 6.16 (d, 1H, $J_{1',2'}$ = 3.0 Hz, H-1'); 5.79 (d, 1H, $J$ = 8.5 Hz, NH-Ile); 5.34 (dd, 1H, $J_{2',3'}$ = 6.1 Hz, $J_{2',1'}$ = 3.0 Hz, H-2'); 5.02-4.96 (m, 2H, H-3', H-4'); 4.41-4.38 (m, 1H, H-5a'); 4.05-3.97 (app t, 1H, $J_{5b',5a'}$ = 4.2 Hz H-5b'); 3.66 (dd, 1H, $J$ = 8.4 Hz, $J$ = 5.0 Hz, CHα-Ile); 3.49-3.43 (m, 1H, CH-Ile); 3.18 (br s, 1H, NH-$SO_2$); 2.60 (s, 3H, $CH_3$-Ile); 1.70-1.62 (m, 2H, $CH_2$-Ile); 1.55 (s, 3H, $CH_3$-Ile); 1.38 (s, 3H, $CH_3$); 1.36 (s, 9H, 3×$CH_3$-Boc); 1.32 (s, 3H, $CH_3$). ESI-MS: *m*/*z* $[M+H]^+$ calc for $C_{24}H_{37}N_7O_9S$: 599.2, found: 600.5.**

#### Compound 3 $^1$H NMR spectrum

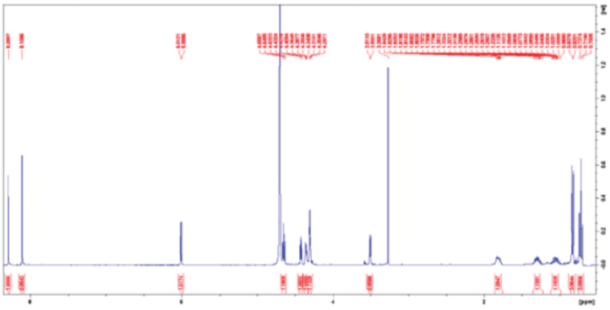

**Yield: 37 mg (60%). White crude foam. $R_f$ = 0.10 ($CHCl_3$ / MeOH, 3:1). $^1$H NMR ($D_2O$) δ/ppm: 8.29 (s, 1H, H-8); 8.11 (s, 1H, H-2); 6.11 (d, 1H, $J_{1',2'}$ = 5.5 Hz, H-1'); 4.65 (app t, 1H, $J$ = 5.3 Hz, H-2'); 4.42 (app t, 1H, $J$ = 4.6 Hz, H-3'); 4.36-4.34 (m, 1H, H-4'); 4.31-4.30 (m, 2H, H-5a', H-5b'); 3.51 (d, 1H, $J$ = 4.4 Hz, CHα-Ile); 1.84-1.78 (m, 1H, CH-Ile); 1.35-1.23 (m, 1H, $CH_2$-Ile); 1.11-0.99 (m, 1H, $CH_2$-Ile); 0.83 (d, 3H, $J$ = 7.0 Hz, $CH_3$-Ile); 0.72 (t, 3H, $J$ = 7.4 Hz, $CH_3$-Ile). ESI-MS: *m*/*z* $[M+H]^+$ calc for $C_{16}H_{25}N_7O_7S$: 459.15, found: 460.4.**

#### Sulfamoyl chloride IR spectrum

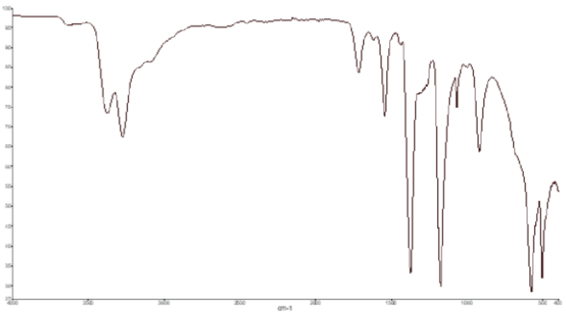

**Wavenumber / $cm^{-1}$:**
**3379.2, 3279.4 (m, NH), 1373.8, 1176.8 (s, S=O)**

**Supplementary Fig. 16: Chemical synthesis of Ile-AMS (compound 3). (A)** Schematic representation of the Ile-AMS synthesis steps. Reagents and conditions are described in Supplementary methods. **(B)** $^1$H NMR spectra for compounds 1-3 and IR spectrum for sulfamoyl chloride.
